# Neuropsychiatric mutations delineate functional brain connectivity dimensions contributing to autism and schizophrenia

**DOI:** 10.1101/862615

**Authors:** Clara Moreau, Sebastian Urchs, Kumar Kuldeep, Pierre Orban, Catherine Schramm, Guillaume Dumas, Aurélie Labbe, Guillaume Huguet, Elise Douard, Pierre-Olivier Quirion, Amy Lin, Leila Kushan, Stephanie Grot, David Luck, Adrianna Mendrek, Stephane Potvin, Emmanuel Stip, Thomas Bourgeron, Alan C. Evans, Carrie E. Bearden, Pierre Bellec, Sebastien Jacquemont, Simons Variation in Individuals Project Consortium

**Author notes:** Shared 1st authorship (Clara Moreau, Sebastian Urchs). Shared senior authorship (Carrie E. Bearden, Pierre Bellec, Sebastien Jacquemont).

## Abstract

16p11.2 and 22q11.2 Copy Number Variants (CNVs) confer high risk for Autism Spectrum Disorder (ASD), schizophrenia (SZ), and Attention-Deficit-Hyperactivity-Disorder (ADHD), but their impact on functional connectivity (FC) remains unclear.

We analyzed resting-state functional magnetic resonance imaging data from 101 CNV carriers, 755 individuals with idiopathic ASD, SZ, or ADHD and 1,072 controls. We used CNV FC-signatures to identify dimensions contributing to complex idiopathic conditions.

CNVs had large mirror effects on FC at the global and regional level. Thalamus, somatomotor, and posterior insula regions played a critical role in dysconnectivity shared across deletions, duplications, idiopathic ASD, SZ but not ADHD. Individuals with higher similarity to deletion FC-signatures exhibited worse cognitive and behavioral symptoms. Deletion similarities identified at the connectivity level could be related to the redundant associations observed genome-wide between gene expression spatial patterns and FC-signatures. Results may explain why many CNVs affect a similar range of neuropsychiatric symptoms.

## Introduction

Copy number variants (CNVs) are deletions or duplications of DNA segments and represent an important source of genetic variation. An increase in rare CNV burden has been linked to a range of neurodevelopmental and psychiatric conditions ^1,2^. Twelve recurrent CNVs have been individually associated with autism spectrum disorder (ASD) ^3^, eight with schizophrenia (SZ) ^4^, and eight with attention deficit hyperactivity disorder (ADHD) ^5^ but the mechanisms by which they lead to neuropsychiatric disorders remain unclear. Although they have large impacts on neurodevelopment, their effect alone does not lead to a psychiatric diagnosis. CNVs could, therefore, be leveraged to identify major dimensions contributing to complex idiopathic conditions.

CNVs at the proximal 16p11.2 and 22q11.2 genomic loci are among the most frequent large effect-size genomic variants and alter the dosage of 29 and 50 genes, respectively ^6,7^. They confer high risk for ASD (10-fold increase for the 16p11.2 deletion and duplication) ^3^, SZ (> 10-fold increase for the 22q11.2 deletion and 16p11.2 duplication) ^4^, and ADHD ^8–12^. Gene dosage (deletions and duplications) affect the same neuroimaging measures in opposite directions (mirror effect). Structural alterations of the cingulate, insula, precuneus and superior temporal gyrus overlap with those observed in meta-analytical maps of idiopathic psychiatric conditions including ASD and SZ.^10,11^

Large effect-size mutations can shed light on pathways connecting genetic risk to brain endophenotypes, such as functional connectivity (FC). FC represents the intrinsic low-frequency synchronization between different neuroanatomical regions. It is measured by means of resting-state functional magnetic resonance imaging (rs-fMRI) which captures fluctuations of blood oxygenation as an indirect measure of neural activity across brain areas when no explicit task is performed ^13,14^. Robust functional brain networks measured by rs-fMRI are also recapitulated by spatial patterns of gene expression in the adult brain ^15,16^.

Few studies have investigated the effect of ‘neuropsychiatric’ CNVs on FC. Dysconnectivity of thalamic-hippocampal circuitry ^17^ has been reported in 22q11.2 deletion carriers, with prominent under-connectivity of the default mode network (DMN), which was predictive of prodromal psychotic symptomatology ^18,19^. Impaired connectivity of long-range connections within the DMN has also been reported by other studies ^20^. A single 16p11.2 study has shown a decrease in connectivity of frontotemporal and -parietal connections in deletion carriers ^21^. These initial studies have focussed on regions of interest but connectome-wide association studies (CWAS) analysing all connections without *a priori* hypotheses have not yet been performed in CNV carriers. Furthermore, their relation to idiopathic conditions has not been investigated.

Brain intermediate phenotypes of psychiatric conditions have mainly been studied by adopting a *top-down* approach, starting with a clinical diagnosis and moving to underlying neural substrates and further down to genetic factors ^22^. Studies applying this analytical strategy in ASD have repeatedly shown patterns of widespread under-connectivity with the exception of overconnectivity in cortico-subcortical connections, particularly involving the thalamus ^23,24^. SZ also exhibits a general under-connectivity profile, mainly involving the medial prefrontal cortex, the cingulate and the temporal lobe ^25^, with over-connectivity of the thalamus ^26^. These altered networks do not appear to be disorder-specific and have been reported across several disorders, including ASD, ADHD, and SZ ^27^. These similarities seem to be distributed across several continuous dimensions ^28^ which may be related to shared genetic contribution across diagnoses, which is documented for common ^29^ and rare ^30^ variants, including the 16p11.2 and 22q11.2 CNVs.

We posit that seemingly distinct genetic variants and idiopathic disorders have overlapping patterns of dysconnectivity, which may help identify FC dimensions, providing insight into the complex connectivity architecture involved in psychiatric conditions.

We aimed to 1) characterize the FC-signatures of four high-risk neurodevelopmental CNVs, 2) explore whether FC-signatures of CNVs represent dimensions observed in idiopathic ASD, SZ, or ADHD and 3) investigate the relationship between deletions at the FC and gene expression level.

To this end, we performed CWAS studies on 101 carriers of a 16p11.2 or 22q11.2 CNV, 122 of their respective controls, 751 individuals with idiopathic ASD, SZ, or ADHD and 948 of their respective controls. To our knowledge, this is the first connectome-wide study to compare rare genomic variants and idiopathic psychiatric conditions.

## Results

### 16p11.2 and 22q11.2 CNVs have large effects on functional connectivity at the global and regional level

The 16p11.2 deletion showed a global increase in FC compared to controls with a mean shift = 0.29 z-scores (p=0.048, permutation test, Figure 1a,c; Supplementary Table S1.1). We observed 88 significantly altered connections (FDR, q < 0.05), and all but one were overconnected with beta values ranging from 0.76 to 1.34 z-scores. Overconnectivity predominantly involved the frontoparietal, somatomotor, ventral attention, and basal ganglia networks (Figure 1c). Regions showing the strongest mean connectivity alterations included the caudate nucleus, putamen, lateral frontal pole, anterior middle frontal gyrus, and dorsal anterior cingulate cortex (Supplementary Table S1.8).

**Figure 1.**
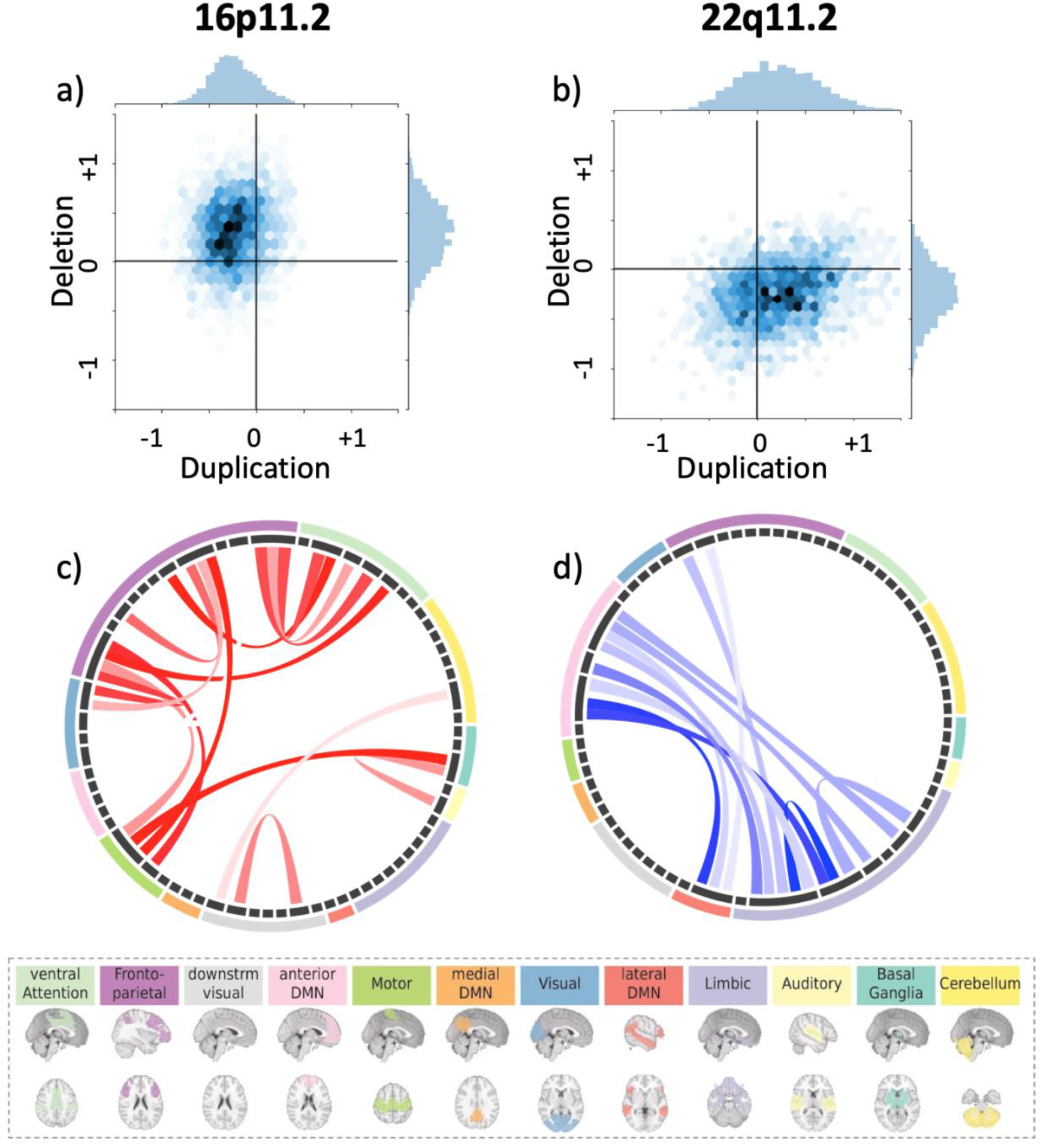
Connectome-wide effects of CNVs. (a-b): Scatterplot (*hexagonal plot*), showing estimates (beta values) from connectome-wide association studies (CWAS) performed between 16p11.2 (a) and 22q11.2 (b) CNVs and their respective controls. In total, 2,080 beta estimates were obtained from a linear model computed from z-scored connectomes based on the variance of the respective controls. The color hue represents the number of beta estimates in the hexagon bin. Y-axis: beta values associated with deletions (CWAS comparing deletions *vs* controls). X-axis: beta-values associated with duplications (CWAS comparing duplications *vs* controls). (c-d): Each chord diagram shows the top 20% of connections surviving FDR correction (*q* < 0.05) from the 16p11.2 deletion (c) and 22q11.2 deletion (d) CWAS. Each chord represents a significantly altered connection between two functional seed regions. All 64 seed regions are represented in the dark grey inner circle. The width of the seed region in the grey inner circle corresponds to the number of altered connections. Seed regions are grouped into 12 functional networks (outer ring, Supplementary Table S1.9). Networks are represented in 12 brains below the two diagrams. Red chords represent overconnectivity and blue chords underconnectivity.

The 22q11.2 deletion was associated with a non-significant global decrease in connectivity Supplementary Table S1.3), with 68 negative connections surviving FDR correction (beta values ranging from −0.68 to - 1.64 z-scores, Figure 1b, d) when compared with control subjects. Underconnectivity predominantly involved the anterior and lateral DMN, and limbic network (Figure 1d). The temporal pole, the ventral anterior insula and peri-insular sulcus, the amygdala-hippocampal complex, the dorsal anterior cingulate cortex, and perigenual anterior cingulate cortex showed the strongest changes in connectivity (see Supplementary Table S1.8).

For 16p11.2 duplication carriers, none of the individual connections survived FDR correction (Figure 1a and Supplementary Table S1.2) relative to controls and none of the individual connections survived FDR correction. For the 22q11.2 duplications, only 16 connections survived FDR and these included overconnectivity in the posterior medial and lateral visual network, the cerebellum I-V, and the lateral fusiform gyrus (Figure 1c and Supplementary Table S1.4 and S1.8).

Deletions and duplications at both loci showed a mirror effect at the global connectivity level. 16p11.2 deletions and duplications also showed mirror effects at the network level (p = 0.006, two-sided). This was not the case for 22q11.2 (Supplementary Results).

A sensitivity analysis showed that results are unaffected by differences in age distribution between deletions and control groups as well as the number of remaining frames available after scrubbing (see Supplementary Results).

### The effect sizes of deletions and duplications are twice as large as the effects of idiopathic SZ, ASD, or ADHD

We performed three independent CWAS, comparing FC between patients with ASD, SZ, ADHD, and their respective controls. Idiopathic SZ showed overall underconnectivity affecting 835 connections, in line with previous reports ^26,31^ (Figures 3a, 3c, Supplemental Results and Supplemental Figure 4, Tables S1.6 and S1.8). Over-connectivity was restricted to 24 connections (FDR, *q* < 0.05).

Idiopathic ASD also showed overall underconnectivity (73 under and 2 overconnected survived FDR, *q* < 0.05, Figure 3b, 3c, Supplemental Results and Supplemental Figure 4, Tables S1.5 and S1.8).

For ADHD, none of the individual connections survived FDR correction (Supplemental Results and Tables S1.7 and S1.8). Sensitivity analyses excluding females from the SZ and ADHD cohorts showed identical results (Supplementary Results).

Among idiopathic conditions, the effect size of connectivity alteration was the highest in SZ (largest beta value = −0.56 std of the control group), followed by autism (largest beta value = −0.46), and ADHD (largest beta value = +0.26). Effect sizes observed for both deletions were approximately two-fold larger (beta values = +1.34 and −1.64 for 16p11.2 and 22q11.2 respectively) than those observed in idiopathic SZ, ASD, and ADHD (Figure 3c). The largest effect size among the 16 connections surviving FDR for the 22q11.2 duplication was Cohen’s d = 1.87.

### Individuals with ASD and SZ relative to controls, show similarities with whole-brain FC-signatures of CNVs

We tested the spatial similarity between whole-brain FC-signatures across CNVs and idiopathic psychiatric conditions. To this mean we computed the similarity (Pearson R) between group-level FC-signatures and the individual connectomes of either cases or controls from another group (Figure 2). This was repeated 42 times between all CNVs and conditions and in both directions. Most of the significant whole-brain FC similarities were observed between individuals with either idiopathic ASD, SZ and 4 CNVs (Figure 3d). ADHD did not show any significant similarities with any other group.

**Figure 2:**
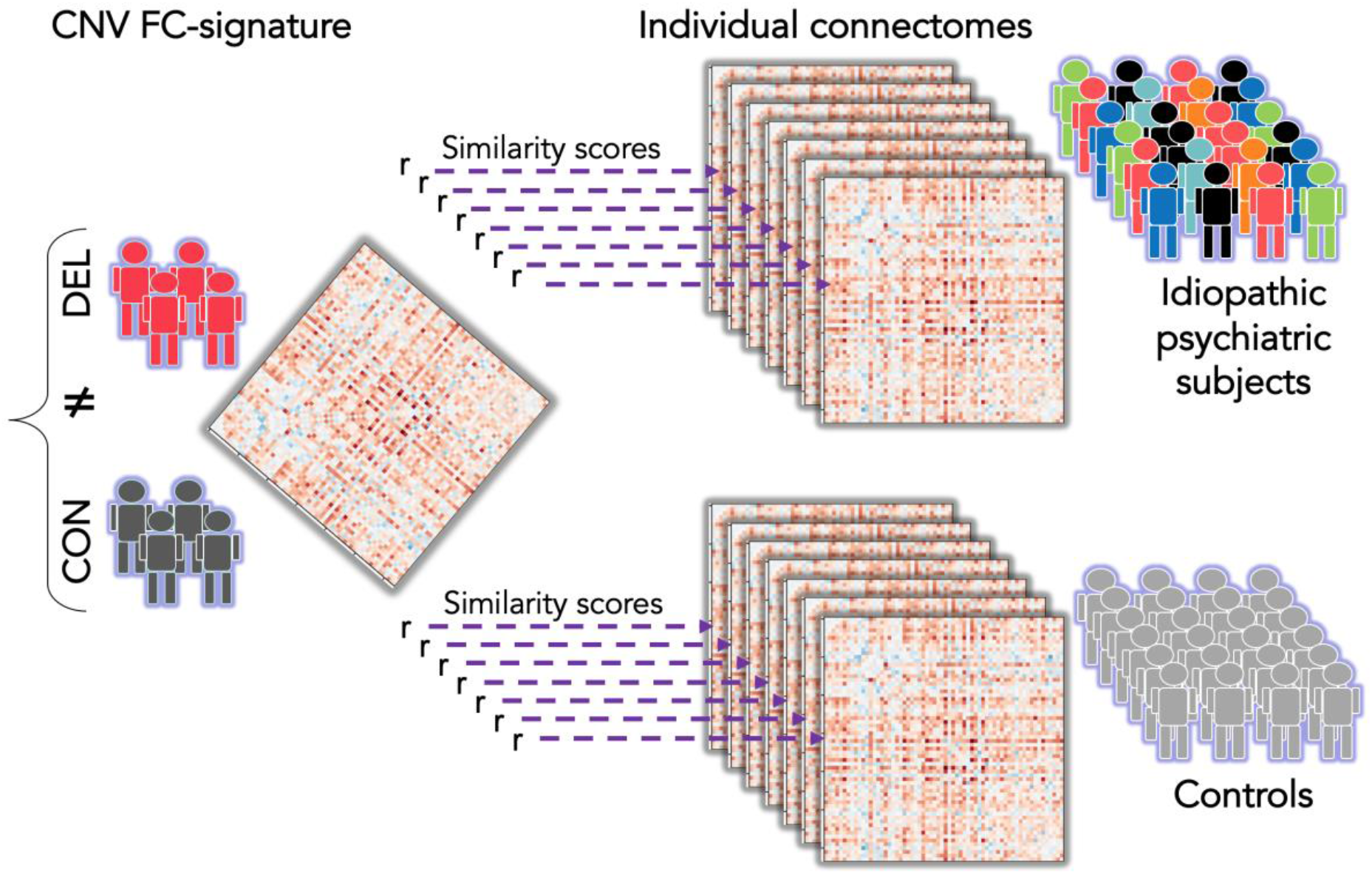
Testing similarities across CNVs and idiopathic conditions. Similarities of FC-signatures across this study were characterized by correlating (Pearson’s r) a group level FC-signature with individual connectomes from either cases and controls. The r values obtained for all cases and all controls were compared using a Mann Whitney test. Here, the group level connectome is represented by a matrix of 2080 beta values, on the left side. It is obtained by contrasting deletion cases (red) and controls (dark grey). The beta map is correlated to 7 individual connectomes of psychiatric cases and 7 connectomes of controls. The different colors used for psychiatric cases represent phenotypic heterogeneity.

**Figure 3.**
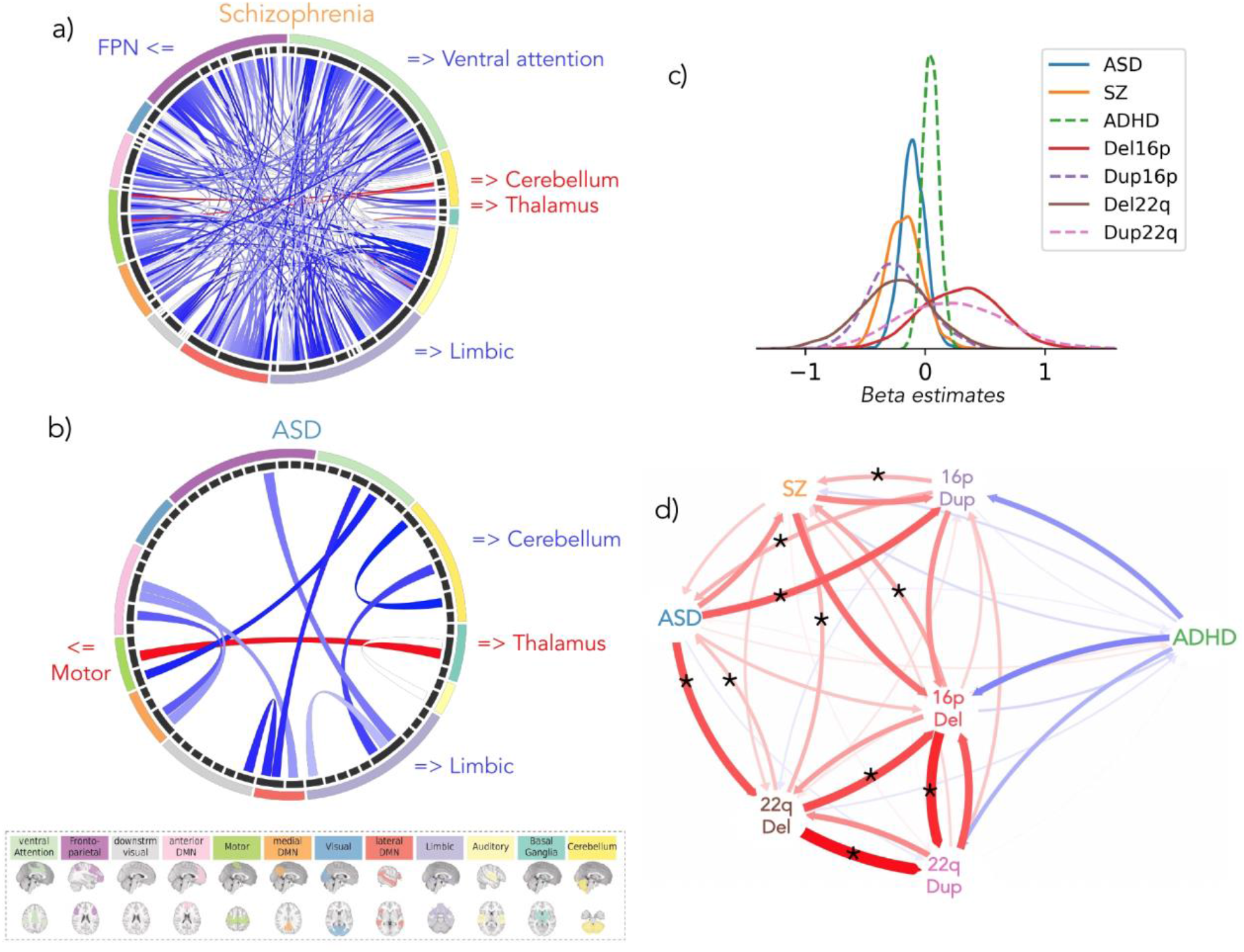
Similarities at the connectome-wide level across ASD, SZ, and deletion FC-signatures. (a, b): Each chord diagram shows the top 20% connections surviving FDR correction (*q* < 0.05) from the SZ (a) and ASD (b) CWAS. Each chord represents a significantly altered connection between two functional seed regions. All 64 seed regions are represented in the dark grey inner circle. The width of the seed region in the grey inner circle corresponds to the number of altered connections. Seed regions are grouped into 12 functional networks (outer ring, Supplementary Table S1.9). The network colors correspond to the legend below. Red chords represent overconnectivity and blue chords underconnectivity. (c) Density plots represent the distribution of 2080 beta estimates for the CWAS (whole brain contrast of cases versus controls) for the SZ, ASD, ADHD, deletion and duplication groups. X-axis values = z-scores of Beta estimates, which were obtained from linear models computed using z-scored connectomes based on the variance of the respective controls. (d) The spatial similarity of whole-brain FC-signatures between CNVs and idiopathic psychiatric conditions. Arrows represent the correlation between group-level FC-signatures and the individual connectomes of either cases or controls from another group. The correlation was computed in both directions. Red and blue arrows represent positive and negative correlations respectively. Arrow thickness represents the effect size of the Mann-Whitney test. Stars represent similarities (Mann-Whitney tests) surviving FDR. ASD: autism spectrum disorder; SZ: schizophrenia; ADHD: attention deficit hyperactivity disorder; FPN: fronto-parietal network; 16pDel: 16p11.2 deletion; 22qDel: 22q11.2 deletion, 16pDup: 16p11.2 duplication; 22qDup: 22q11.2 duplication

### Thalamus and somatomotor regions play a critical role in dysconnectivity observed across CNVs and idiopathic psychiatric conditions

We asked if whole-brain FC similarities between individuals with ASD, SZ and CNVs may be driven by particular regions. We thus repeated the same similarity analysis presented above at the level of the FC signatures of each of the 64 seed regions. Individuals with SZ showed increased similarity with 28 out of the 64 regional FC-signatures of the 16p11.2 deletion than controls (FDR, *q* < 0.05). They also showed increased similarity with 18 region-level FC-signatures of the 22q11.2 deletion (Figure 4, Supplementary Tables S2.3 and S2.4, and Results). Deletion FC-signatures did not show any similarity with controls. We ranked the effect size of each seed region and compared them for both deletions. The seed regions with the highest similarity between SZ and 16p11.2 were also those with the highest similarity between SZ and 22q11.2 (adjusted R2 = 0.37, p = 6e-08) (see Supplementary Results). Sensitivity analysis showed that the same regions are driving similarities with SZ and ASD irrespective of psychiatric diagnoses in 22q11.2 deletion carriers (see Supplementary Results).

**Figure 4.**
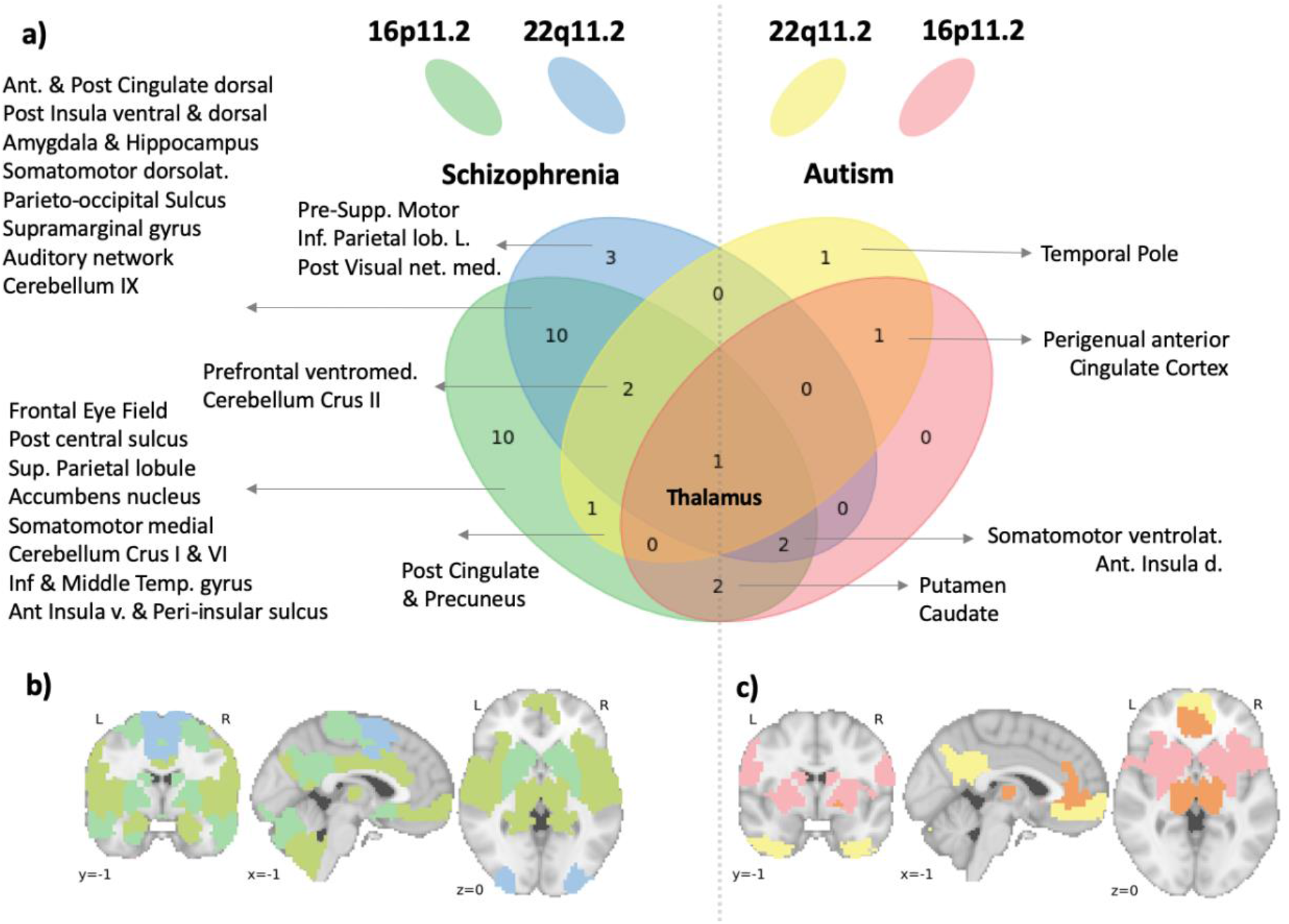
Regional similarity between the individual FC profiles of subjects with a psychiatric diagnosis and FC-signatures of 16p11.2 and 22q11.2 deletions. The FC-signatures of both deletions are decomposed into 64 seed-regions. Deletion FC-signatures are correlated to the individual connectivity profile of subjects with a psychiatric diagnosis and their respective control subjects. Of note, the correlation is equivalent to the mean centring of all region-based FC-signatures. Significantly higher similarities of patients with either ASD and SZ were present in 33 seeds regions (FDR) and are presented on the right and the left side of the Venn diagram, respectively (a), and also in the corresponding left (b) and right (c) brain maps. At the intersection of all ellipses, the thalamus FC-signatures of both deletions showed increased similarity with individuals who have a diagnosis of ASD or SZ compared to their respective controls. 16pDel: 16p11.2 deletion, 22qDel: 22q11.2 deletion; Ant: anterior; Post: posterior; dorsolat: dorsolateral; Inf: inferior; L.: left; v.: ventral; net.: network; med: medial; Supp: supplementary; lob: lobule (Full-name labels are provided in Supplementary Table S1.9).

Individuals with autism showed greater similarity with six regional FC-signatures of the 16p11.2 deletion compared to controls (FDR, *q* < 0.05). They also showed greater similarity with six region-level FC-signatures of the 22q11.2 deletion (Figure 4, Supplementary Tables S2.1 and 2.2, and Results). Deletion FC-signatures did not show significant similarities with controls for any of the 64 seed regions. Of note, individuals with SZ and ASD showed higher similarity with the thalamus FC-signatures of both deletions (Figure 4). None of the similarities correlated with motion or sex. Regions driving similarities between psychiatric conditions and deletion FC signatures were also those with the highest number of connections altered by each deletion individually. Eight and six out of the top 10 regions altered by 22q11.2 and 16p11.2 respectively were driving similarities with psychiatric conditions (Supplementary Table 1.8).

Despite lower power, we investigated similarities with duplication FC-signatures. The number of significant regional similarities was smaller. Out of the 28 regions showing a similarity between idiopathic conditions and duplications, 17 regions also showed similarities with deletions (See Supplementary Results). Individuals with ADHD did not show higher similarities with the regional FC-signatures of any CNVs except for 2 regional FC-signatures of the 16p11.2 duplication (Supplementary Tables S2.5 and S2.6).

### Similarity to deletion FC-signatures is associated with symptom severity

We investigated whether regional FC similarities with deletions described above are associated with symptom severity among individuals in idiopathic psychiatric cohorts. Symptom severity was assessed using the Autism Diagnostic Observation Schedule (ADOS, ^32^), in ASD, Positive and Negative Syndrome Scale (PANSS, ^33^) in SZ, and Full Scale Intelligence Quotient (FSIQ) in ASD. The 10 seed regions with significant FC similarity between ASD and either deletion were those showing the strongest association with the ADOS symptom-severity score (two regions passed FDR correction *q* < 0.05: the caudate nucleus and temporal pole) and FSIQ (Figure 6 and Supplementary Tables S3.1-3.4). Among the seed regions contributing to the similarity between SZ individuals and deletions, none were significantly associated with PANSS measures after FDR correction. FSIQ data was not available in the SZ cohorts.

### 16p11.2 and 22q11.2 deletions show regional FC similarities

Although the two deletions showed opposing effects on global connectivity (Figure 3c), their connectomes were positively correlated (Figure 3.d). We, therefore, sought to identify the main regions that contributed to this connectome-wide correlation.

Using the same approach as above (Figure 2), we correlated the 22q11.2 deletion group-level FC-signature with individual connectomes of 16p11.2 deletion carriers and their respective controls. The 22q11.2 deletion FC-signature showed significant similarities with 16p11.2 deletion carriers for 12 regions (FDR, q<0.05), mainly involving the frontoparietal, ventral attentional, and somatomotor networks. The reverse test showed significant similarity of the 16p11.2 deletion FC-signature with 22q11.2 deletion carriers in 10 regions within the anterior and lateral DMN, frontoparietal and basal ganglia networks. Four seed regions were observed in both tests (Figure 5a). We reasoned that the FC-similarity between CNVs may be informed by the spatial patterns of gene expression within both genomic intervals.

**Figure 5.**
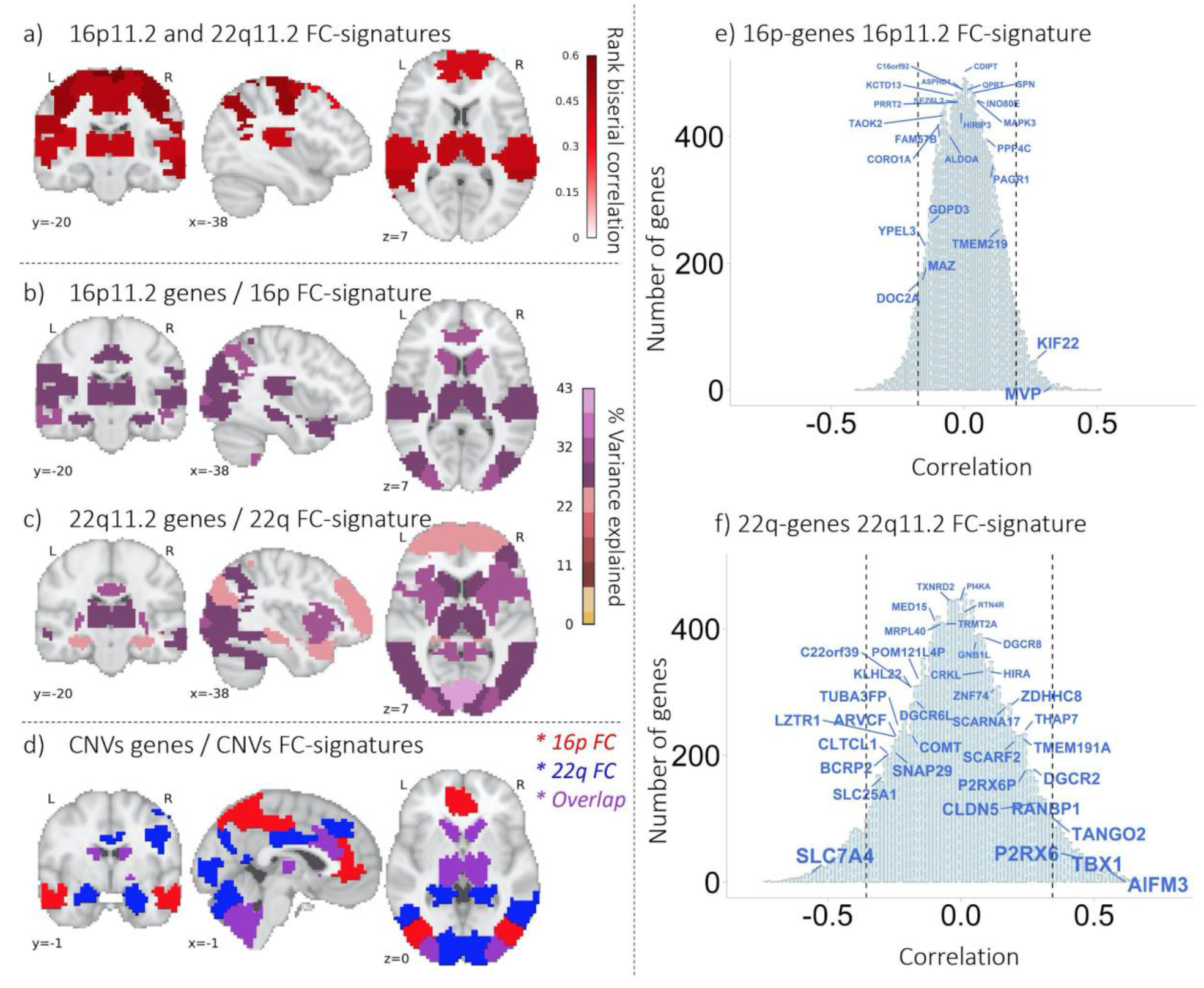
FC similarities between 16p11.2 and 22q11.2 and relationship with gene expression. Legend: (a) FC similarities between both deletions at the regional level. The values in the brain map represent the level of the FC similarity between deletions (rank biserial correlation, Mann Whitney test). The values are thresholded (FDR, 64 regions): 18 out of 64 regions are similar between deletions. (b) Relationship between spatial patterns of gene expression within the 16p11.2 locus and regional FC signatures of the 16p11.2 deletion. A partial least square regression (PLSR) was conducted for each of the 64 regions. Maps are thresholded (FDR corrected for 64 regions) and color code represents the percentage of variance explained by gene expression using 2 components in the PLSR. c) The same analysis was conducted for 22q11.2 genes and the 22q11.2 deletion FC signature. (b,c) Eleven regions overlapped across PLSR maps: thalamus, caudate, anterior insula and posterior insula sulcus, amygdala and hippocampus, cerebellum 9 and right crus-2, dorsal anterior cingulate, left inferior parietal lobule, medial posterior visual network, lateral posterior visual network, and dorsal visual network. Three (over the 18) regions identified in the between deletion similarity (a) analysis are also present in the gene expression/FC-signature association maps (b,c): Thalamus, dorsal anterior cingulate and left inferior parietal lobule. (d) Low specificity for the relationship between spatial patterns of gene expression and regional FC deletion signatures. In red: the 16p11.2 regional FC associated with the expression patterns of both the 16p11.2 and the 22q11.2 genes. In blue, the 22q11.2 regional FC is associated with the expression patterns of genes in both genomic loci. In purple, 7 regions were found with both deletion FC-signatures, and the expression patterns of genes encompassed in both genomic loci. (e-f) Expression patterns of genes within and outside CNVs correlate with FC-signatures of 16p11.2 and 22q11.2 deletions. The light blue histogram represents the distribution of correlations for 15663 genes with available gene expression data from the AHBA. X-axis values : Pearson coefficients. Y-axis values: number of genes. Genes within the CNVs have font-size scaled based on p values. Dotted lines represent the 5th and 95th percentiles of the correlation distribution genome-wide.

**Figure 6.**
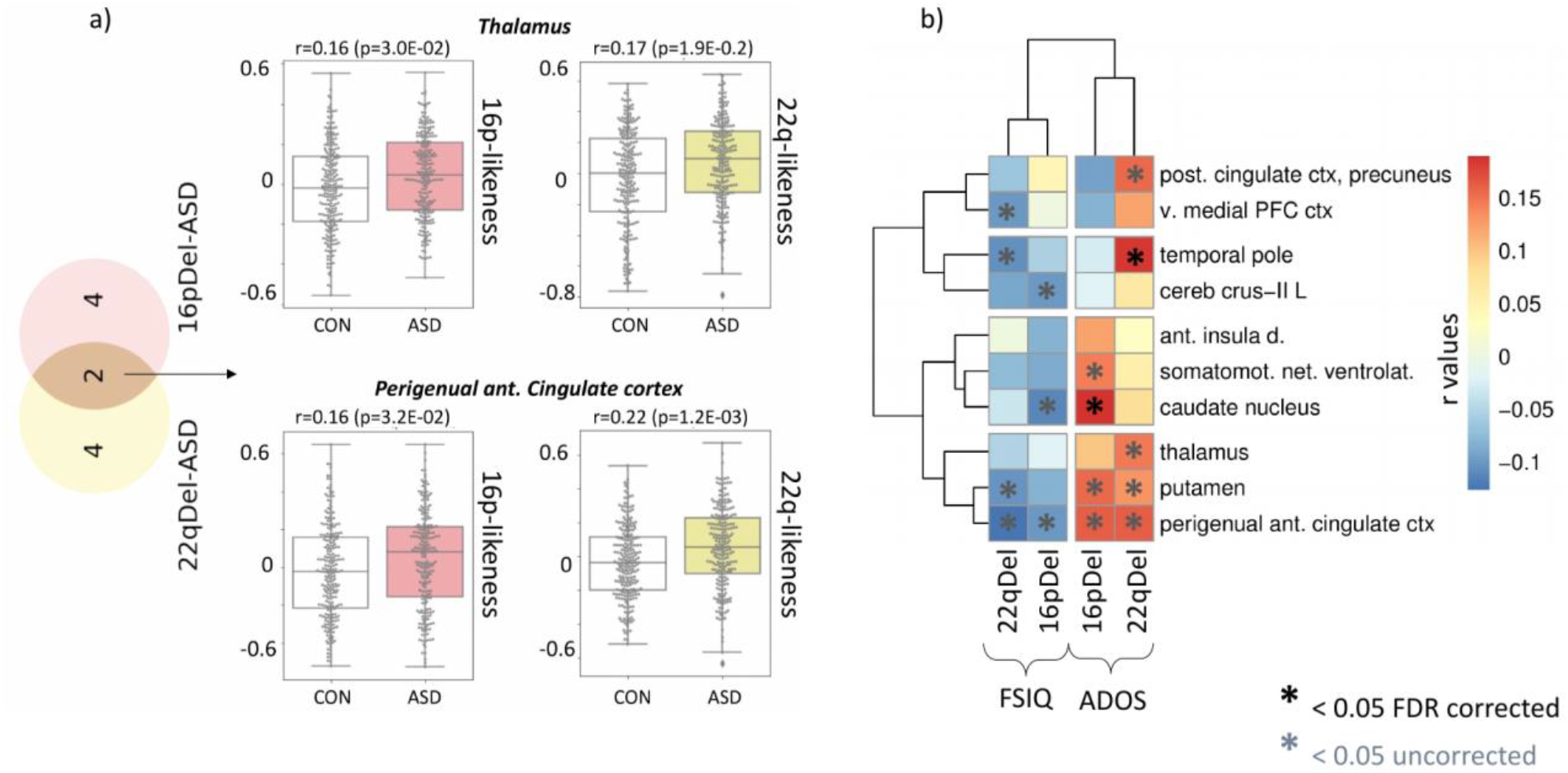
Relationship between the deletion FC-signatures and behavior. a) Boxplots represent the connectivity similarity for two seed regions (thalamus and perigenual anterior cingulate cortex). Each data point represents one individual: r-value of the Pearson correlation between the deletion FC-signatures and the FC-profile of an individual with ASD (n = 221 in the coloured boxplots) or a control subject (n = 232 in the non-coloured boxplots). For the two seed regions, individuals with ASD show significantly higher similarity (FDR, *q* < 0.05) with the 16p11.2 (pink) and 22q11.2 (yellow) deletion FC-signatures than the individual controls. All seed regions showing significantly higher similarity with ASD are represented in the Venn diagram on the left. b) We investigated the relationship with cognitive scores and found that stronger individual similarity with the deletion FC-signature was associated with more severe symptoms measured by FSIQ and ADOS. Heatmaps show the level of correlation between behaviour scores and the similarity with deletion FC-signatures 16pDel: 16p11.2 deletion, 22qDel: 22q11.2 deletion; FSIQ: full-scale intelligence quotient; ADOS: autism diagnostic observation schedule; ant.: anterior; post: posterior; v.: ventral; PFC: prefrontal cortex; cereb: cerebellum; d.: dorsal; L: left; ctx: cortex; net.: network. (Full-name labels are provided in Supplementary Table S1.9).

### Association between gene expression spatial patterns and deletion FC-signatures

We performed Partial Least Squares Regression (PLSR) to investigate the association between FC-signatures of each deletion and the expression patterns of 37 and 24 genes encompassed in the 22q11.2 and 16p11.2 genomic loci respectively. The 2 components required to reach a significant association explained 24.2% of the variance of the 16p11.2 deletion FC profile (p=0.041, 5000 random FC profiles). For the 22q11.2 deletion, either one or 2 components were significant (p<0.0002, 5000 random FC profiles). The 2 components explained 43.2% of the variance of the 22q11.2 deletion FC signature. Similar PLSR analyses performed for each of the 64 regions showed that 18 and 32 regional FC-signatures were significantly associated (5000 random FC-signatures, FDR 64 regions) with spatial patterns of gene expression at the 16p11.2 and 22q11.2 loci respectively (Figure 5b-c). However, this relationship was not specific because 22q11.2 genes were also associated with n=20 regions of the 16p11.2 FC signature. Conversely, the 16p11.2 genes were associated with n=19 regions of the 22q11.2 FC signature (Figure 5.c, Supplementary Table S4.2).

To further investigate the low specificity of the connectivity/gene expression relationship, we tested the individual correlation (Pearson) of all 15663 genes with available expression data with deletion FC signatures (Figure 5d, e, Supplementary Table S4.3-S4.4). Correlations with the 16p11.2 and 22q11.2 FC signatures were observed for 421 and 3883 genes respectively (5000 random FC-signatures). After genome-wide FDR correction, the expression of 1834 genes remained spatially correlated with 22q11.2 and none with the 16p11.2 deletion FC signature. The median correlation values for the n=24 16p11.2 genes and the 16p11.2 FC signature was not higher than the median correlation of 10000 randomly sampled gene sets of the same size (n=24, p=0.31). The same was true for 22q11.2 genes (n=37, p=0.36). However, both deletions were enriched in genes with correlations ranking higher than the genome-wide 98th percentile: *MVP* and *KIF22* showed correlations (r_*MVP*_=0.33, r_*KIF22*_=0.26, Supplementary Table S4.4) ranking at the 99.76th and 98.76th percentile genome-wide (p=0.03; null:10000 random gene sets). For 22q11.2, *AIFM3, TBX1* and *P2RX6* showed correlations in the 99.5, 98.84, 98.33th percentile (p=0.04).

## Discussion

This proof of concept study provides the first connectome-wide characterization of four CNVs that confer high risk for psychiatric disorders. Deletions and duplications at the 16p11.2 and, to a lesser extent, the 22q11.2 locus were associated with mirror effects at the connectome-wide level. Overconnectivity in the 16p11.2 deletion predominantly involved the ventral attention, motor, and frontoparietal networks. Underconnectivity in the 22q11.2 deletion involved the anterior and lateral DMN and the limbic network. Regional FC-signatures of deletions and duplications, in particular, those implicating the thalamus, somatomotor, posterior insula and cingulate showed significant similarities with the complex architecture of idiopathic ASD, SZ but not ADHD. Seemingly distinct, rare neuropsychiatric mutations may converge on dimensions representing mechanistic building blocks shared across idiopathic conditions. The spatial expression patterns of genes encompassed in both genomic loci were associated with FC-signatures of the corresponding deletion but many genes outside these 2 loci also show similar levels of association. This redundancy may represent a factor underlying shared FC signatures between both deletions.

22q11.2 and 16p11.2 CNVs showed large effect sizes on FC that are similar to those previously reported for structural neuroimaging measures, cognition, and behaviour ^8,10,11^. In sharp contrast, there is significant discordance between the severe clinical manifestations observed in idiopathic ASD and SZ, and the small effect size observed in case-control studies at the FC level. Previous structural neuroimaging studies of the same idiopathic psychiatric conditions have also reported small effect sizes ^34,35^. This discordance may be due to the heterogeneity of these idiopathic conditions and hints at the presence of subgroups or latent dimensions associated with larger effect sizes ^28^.

The FC-signatures of both deletions (and to a lesser extent those of both duplications) showed similarities with autism and schizophrenia, but not ADHD. Regions contributing to these similarities were also those with the highest number of connections altered by each deletion individually. The FC-signature of the same seed regions also showed the highest association with ASD severity scores and general intelligence in the idiopathic autism sample.

Among the regions, results highlighted overconnectivity between the thalamus and sensory-motor, auditory and visual networks as a common alteration across CNVs and individuals with idiopathic autism or schizophrenia who do not carry CNVs. This is in line with recent rs-fMRI studies performed across psychiatric illnesses ^28^. Perceptual dysfunctions are core features of SZ and ASD ^36^. Those include auditory and visuals hallucinations in SZ ^37,38^, impairments in gestalt visual perception and discrimination of visual motion ^39^, disturbances in auditory and tactile discrimination in Autism. Impairments in phonology ^40^, as well as visual and auditory deficits, have also been demonstrated in 16p11.2 and 22q11.2 deletion carriers ^41–43^. A general thalamo-sensory disturbance may, therefore, be central across genomic mutations and psychiatric diagnoses. Further studies are required to investigate genome-wide, the genetic determinants of thalamo-sensory disturbance. Because it appears ubiquitous across conditions, the genetic basis is likely to be very broad. FC similarities between idiopathic psychiatric disorders, deletions and duplications are also in line with an emerging body of literature that points to common neurobiological substrates for mental illness ^44^. Evidence includes the genetic correlation between psychiatric disorders ^29,45^ and pleiotropic effects of CNVs associated with several conditions ^2,46^.

Recent work has shown that many genes share similar spatial patterns of expression 15 organized along broad spatial gradients in the brain that are closely related to functional connectivity networks ^47,48^. In line with these observations, we show that FC-signatures of deletions are associated with expression patterns of genes within as well as outside the genomic loci of interest. The FC profile of the thalamus, dorsal anterior cingulate, and left inferior parietal lobule were associated with expression patterns of genes at both loci and may explain, in part, the FC similarities between both deletions.

Functional connectivity studies using a top-down case-control approach (eg. autism versus control) have characterized large-scale brain network changes associated with diseases, but this framework is unable to describe the directionality of this relationship ^49^. FC-changes may not necessarily represent an intermediate brain phenotype but rather a secondary impact of psychiatric illnesses. Our strategy integrating top-down and bottom-up approaches shows that individuals with idiopathic ASD or SZ as well as CNV carriers who do not meet diagnostic criteria for these conditions share regional FC alterations. This suggests that the risk conferred by genetic variants and the associated FC-patterns represent important dimensions that are necessary but insufficient to cause disease. Additional factors and associated FC-patterns are required (incomplete penetrance ^50^). Bottom-up approaches studying rare variants have almost exclusively been performed individually. Our results suggest, however, that they likely converge on overlapping intermediate brain phenotypes, consistent with a recent study showing overlapping effects on subcortical structures across 12 different CNVs conferring risk to SZ ^51^.

### Limitations

Reproducibility of rs-fMRI in psychiatry has been challenging. However, when studies using similar analytical strategies are compared, there are consistent results. In SZ and ASD, global decrease in FC has been reported by most studies except for those adjusting for global signal ^24,52^. Increased thalamocortical connectivity is also repeatedly reported in both conditions ^24,26,31^. These previous findings are consistent with our results (see Supplementary Results). The 22q11.2 deletion FC signature is consistent with previous works on 22q11.2 FC alterations that showed 1) underconnectivity of the DMN ^53,54^, 2) thalamocortical overconnectivity and underconnectivity involving the hippocampus ^17^. The only rs-fMRI study previously published for the 16p11.2 deletion focused on the dmPFC ^21^. Using the same approach and regressing global signal, we also found underconnectivity of the dmPFC with the same set of regions. This highlights the fact that many seemingly discrepant results can be reconciled once methodologies are aligned.

There is no available genetic data for any of the three idiopathic cohorts. However, the frequency of 16p11.2 and 22q11.2 CNVs in ASD or SZ is < 1% ^3,4^. This suggests that the observed FC similarities between CNVs and ASD or SZ are driven by other factors.

Expression data were derived from 6 adult brains of the AHBA and results should be interpreted with caution.

The results on duplications should be interpreted with caution due to our limited power to detect changes in connectivity. The limited phenotypic data in the SZ group did not allow to investigate the relationship between deletion FC-signatures and cognitive traits in this sample. Lack of similarity observed for ADHD is line with the small association between 16p11.2 CNVs and ADHD but is discordant with the association reported for 22q11.2 ^5^. ADHD has a smaller effect size than SZ and ASD, which may have limited our analysis ^55^. Several confounding factors may have influenced some of the results. Those include sex bias, which is present across all 3 psychiatric cohorts, age differences in the 16p11.2 deletion group, diagnosis of ASD and ADHD in 22q11.2 deletion carriers, and medication status in the idiopathic ASD and SZ groups. However, carefully conducted sensitivity analyses, investigating all of these confounders did not change any of the results.

### Conclusion

Deletion and duplication at several genomic loci result in mirror effects across many human traits ^56–59^, including brain connectivity. Haploinsufficiency may define functional connectivity dimensions that represent building blocks contributing to idiopathic psychiatric conditions. Thalamo-sensory disturbance may represent one dimension central across genomic mutations and psychiatric diagnoses. The redundant associations observed, genome-wide, between gene expression and connectivity may explain similarities across genomic variants and idiopathic conditions and the fact that many CNVs affect a similar range of neuropsychiatric symptoms. It is, therefore, becoming increasingly difficult to justify the study of psychiatric conditions or rare genetic variants in isolation. Large scale studies simultaneously integrating a top-down approach across diagnostic boundaries, and a bottom-up investigation across a broad set of genomic variants are required to improve our understanding of specific and common psychiatric outcomes associated with genetic variants and FC signatures.

## Materials and Methods

### Samples

We performed a series of CWAS using individuals from five data sets (Table 1 and Supplementary Materials and Methods).

**Table 1.** Cohort characteristics. Description of the cohorts after filtering for quality criteria. SVIP: Simons Variation in Individuals Project; UCLA: University of California, Los Angeles; ASD: autism spectrum disorder; ABIDE: Autism Brain Imaging Data Exchange; SZ: schizophrenia; ADHD: attention-deficit/hyperactivity disorder; Del: deletion; Dup: duplication; Age (in years); FSIQ: Full-Scale Intelligence Quotient; M: male; FD: framewise displacement (in mm). Quantitative variables are expressed as the mean ± standard deviation. *More information regarding the remaining number of time frames for each group, and the percentage of motion censoring, is provided in Supplementary Materials and Methods. Sensitivity analyses investigating sex bias in the 3 idiopathic cohorts are presented in Supplementary Results. Sensitivity analysis investigating medication effect in ASD cohort is presented in Supplementary Results. Sensitivity analyses also showed that the FC-signature of 22q11.2 deletions is not influenced by a diagnosis of ASD or ADHD (Supplementary Figure 1). Columns ASD, SZ, and ADHD represent the number of subjects with those diagnoses. One subject may have several diagnoses. For example, 9 subjects with ASD have also an ADHD diagnosis.

| Dataset | Status | n | Age | FSIQ | Sex | FD* | ASD | ADHD | SZ |
| --- | --- | --- | --- | --- | --- | --- | --- | --- | --- |
| **SVIP**<br>**16p11.2 Cohort**<br>**(2 sites)** | Del Carriers | 20 | 12.7 (6.8) | 92.5 (16.1) | 12M | 0.18 (0.03) | 4 | 6 | 0 |
|  | Controls | 79 | 26.7 (14.7) | 103.6 (14.8) | 46M | 0.17 (0.04) | 0 | 0 | 0 |
|  | Dup Carriers | 23 | 28.2 (12.8) | 94.1 (15.1) | 12M | 0.19 (0.05) | 1 | 1 | 0 |
| **UCLA**<br>**22q11.2 Cohort**<br>**(1 site)** | Del Carriers | 46 | 16.8 (6.1) | 77.2 (13.8) | 20M | 0.17 (0.06) | 18 | 20 | 3 |
|  | Controls | 43 | 13.0 (4.6) | 111.9 (17.6) | 22M | 0.14 (0.04) | 0 | 2 | 0 |
|  | Dup Carriers | 12 | 16.74 (13) | 95.7 (19) | 7M | 0.17 (0.1) | 3 | 4 | 0 |
| **ASD Cohort**<br>**(ABIDE 10 sites)** | Cases | 225 | 15.9 (6.5) | 103.7 (17.4) | 221M | 0.18 (0.05) | 225 | - | - |
|  | Controls | 234 | 15.7 (6.1) | 110.63 (12.1) | 232M | 0.17 (0.04) | 0 | - | - |
| **Schizophrenia**<br>**Cohort (10 sites)** | Cases | 241 | 33.62 (9.2) | - | 179M | 0.16 (0.06) | - | - | 241 |
|  | Controls | 242 | 32.3 (9.6) | - | 181M | 0.14 (0.05) | - | - | 0 |
| **ADHD Cohort**<br>**(7 sites)** | Cases | 289 | 11.5 (2.8) | 106.8 (13.7) | 227M | 0.15 (0.04) | - | 289 | - |
|  | Controls | 474 | 12.2 (3.3) | 114..2 (13.1) | 250M | 0.14 (0.04) | - | 0 | - |

1-2) Two genetic-first cohort (recruitment based on the presence of a genetic variant, regardless of any DSM diagnosis):

16p11.2 deletion and duplication carriers (29.6-30.1MB; Hg19), and extrafamilial controls from the Simons Variation in Individuals Project (VIP) consortium ^60^.

22q11.2 deletion and duplication carriers (18.6-21.5MB; Hg19) and extrafamilial controls from the University of California, Los Angeles.

3) Individuals diagnosed with ASD and their respective controls from the ABIDE1 multicenter dataset ^23^.

4) Individuals diagnosed with SZ (either DSM-IV or DSM-5) and their respective controls. We aggregated fMRI data from 10 distinct studies.

5) Individuals diagnosed with ADHD (DSM-IV) and their respective controls from the ADHD-200 dataset^61,62^.

Imaging data were acquired with site-specific MRI sequences. Each cohort used in this study was approved by the research ethics review boards of the respective institutions. Signed informed consent was obtained from all participants or their legal guardian before participation. Secondary analyses of the listed datasets for the purpose of this project were approved by the research ethics review board at Sainte Justine Hospital. After data preprocessing and quality control, we included a total of 1,928 individuals (Table 1).

### Preprocessing and quality control procedures

All datasets were preprocessed using the same parameters with the same Neuroimaging Analysis Kit (NIAK) version 0.12.4, an Octave-based open-source processing and analysis pipeline ^63^. Preprocessed data were visually controlled for quality of the co-registration, head motion, and related artefacts by one rater (Supplementary Materials and Methods).

### Computing connectomes

We segmented the brain into ^64^ functional seed regions defined by the multi-resolution MIST brain parcellation ^64^. FC was computed as the temporal pairwise Pearson’s correlation between the average time series of the 64 seed regions, after Fisher transformation. The connectome of each individual encompassed 2,080 connectivity values: (63×64)/2 = 2016 region-to-region connectivity + 64 within seed region connectivity. We chose the 64 parcel atlas of the multi-resolution MIST parcellation as it falls within the range of network resolution previously identified to be maximally sensitive to functional connectivity alterations in neurodevelopmental disorders such as ASD.^65^

Statistical analyses were performed in Python using the scikit-learn library ^66^. Analyses were visualized in Python and R. Code for all analyses and visualizations is being made available online through the GitHub platform https://github.com/surchs/Neuropsychiatric_CNV_code_supplement.

### Statistical analyses

All of the following analyses are summarised in Supplemental Materials and Methods.

#### Connectome-wide association studies

We performed seven CWAS, comparing Functional Connectivity (FC) between cases and controls for four CNVs (16p11.2 and 22q11.2, either deletion or duplication) and three idiopathic psychiatric cohorts (ASD, SZ, and ADHD). Note that controls were not pooled across cohorts. Within each cohort, FC was standardized (z-scored) based on the variance of the respective control group. CWAS was conducted by linear regression at the connectome level, in which z-scored FC was the dependent variable and clinical status the explanatory variable. Models were adjusted for sex, site, head motion, and age. We determined whether a connection was significantly altered by the clinical status effect by testing whether the β value (regression coefficient associated with the clinical status variable) was significantly different from 0 using a two-tailed t-test. This regression test was applied independently for each of the 2,080 functional connections. We corrected for the number of tests (2,080) using the Benjamini-Hochberg correction for FDR at a threshold of *q* < 0.05 ^67^, following the recommendations of Bellec *et al*. 2015 ^68^.

We defined the global FC shift as the average of the β values across all 2,080 connections and tested for significance using a permutation test. We performed 5000 random CWAS by contrasting CNV carriers and controls after shuffling the genetic status labels. For example, we randomly permuted the clinical status of 16p11.2 deletion carriers and their respective controls in the 16p11.2 deletion *vs* control CWAS. We then estimated the p-value by calculating the frequency of random global FC shifts that were greater than the original observation^69^.

#### Gene dosage mirror effects on functional connectivity

We tested whether networks are affected by gene dosage in a mirror fashion by computing the product of the β values obtained in each genetic group contrasts: “Deletions *vs* Controls” and “Duplications *vs* Controls” (separately for 16p11.2 and 22q11.2). Negative values indicate mirror effects of deletions and duplications on FC. Positive values indicate effects in the same direction for deletions and duplications. The obtained products of the β values were grouped into 12 canonical functional networks using information from the multi-resolution brain parcellation (Supplementary Table S1.9).

#### Similarity of whole-brain FC-signatures between idiopathic psychiatric conditions and CNVs

We tested the similarity between dysconnectivity measured across idiopathic psychiatric conditions and CNV. This similarity was tested by correlating individual whole-brain connectomes of cases and controls of one group to the whole brain FC-signature (group level) of another group (Figure 2). The group-level FC-signature was defined as the 2,080 β values obtained from the contrast of cases vs. controls. This was repeated 21 times between all CNVs and conditions and in both directions (n=42 similarity tests).

Individual connectomes of cases and their respective controls were used after independently adjusting for sex, site, head motion, age, and average group connectivity for each of the datasets.

Similarity scores were derived by computing Pearson’s correlations between the whole brain connectomes. We asked whether cases compared to their respective controls had significantly higher (or lower) similarity to whole-brain FC-signature of another group using a Mann-Whitney U test. We reported significant group differences after FDR correction accounting for the 42 tests (*q* < 0.05).

#### Similarity of regional FC-signatures between idiopathic conditions and CNVs

The same approach described above was performed at the regional level. Each of the 1705 connectomes of individuals with idiopathic psychiatric conditions and their respective controls was independently adjusted for sex, site, head motion, age, and average group connectivity for each dataset. We calculated a similarity score between these individual connectomes and the FC-signatures of the 16p11.2 and 22q11.2 deletions and duplications. The FC-signatures were broken down into 64 region-level FC-signatures and similarity scores were derived by computing Pearson’s correlations between the 64 β values associated with a particular region. For each region, we tested whether individuals with a psychiatric diagnosis had significantly higher (or lower) similarity to 16p11.2 or 22q11.2 deletion FC-signatures than their respective controls using a Mann-Whitney U test. We reported significant group differences after FDR correction (*q* < 0.05) for the number of regions (64).

We investigated the relationship between symptom severity and similarity with deletions. The similarity of individuals with deletion FC-signatures was correlated (Pearson’s r) with cognitive and behavioral measures. Those included the ADOS and FSIQ in the autism sample and the PANSS in the SZ sample. The p-values associated with these correlations were corrected for multiple comparisons (FDR, *q* < 0.05).

#### Similarity between 16p11.2 and 22q11.2 deletions at the regional level

We correlated the 22q11.2 group-level deletion-FC-signature with individual connectomes of 16p11.2 deletion carriers and their respective controls. We correlated as well the 16p11.2 group-level deletion-FC-signature with individual connectomes of 22q11.2 deletion carriers and their respective controls. For each region, we tested whether individuals with a deletion had significantly higher (or lower) similarity to the other deletion FC-signatures than their respective controls using a Mann-Whitney U test. We reported significant group differences after FDR correction (*q* < 0.05) for the number of regions (64).

#### Gene expression analyses

We aligned the gene expression maps from AHBA to the MIST64 functional parcellation following previously published guidelines ^70^ and adapting the abagen toolbox ^71^ (Supplementary methods). For all analyses, we used a dataset including 1 expression value per gene and per functional region. Expression values were associated with the average connectivity alteration of the corresponding regions (mean of all 64 beta values of each region).

PLSR method was used to investigate the association between spatial patterns of gene expression (of the 37 and 24 genes encompassed in the 22q11.2 and 16p11.2 genomic loci) and the 16p11.2 and 22q11.2 FC signatures. PLSR is a multivariate approach, which has previously been applied to investigate the relationship between neuroimaging phenotypes and spatial patterns of gene expression ^72–75^. PLSR was performed separately for 16p11.2 and 22q11.2 genes. Components defined by PLSR were the linear combinations of the weighted gene expression scores (predictor variables) that most strongly correlated with FC-signatures of deletions (response variables). To assess significance, we recomputed PLSR using 5000 null FC-signature maps and counted the number of times the explained variance was higher than the original observation. Null FC-signatures were obtained by computing 5000 times the contrast between CNVs and controls after label shuffling for 16p11.2 and 22q11.2 separately. To investigate the association between FC alterations and expression patterns of individual genes, we computed Pearson correlations. The null distribution was defined by the same 5000 random FC-signatures described above.

To test the specificity of the relationship between gene expression and FC, we randomly sampled 10000 gene sets (n=24 for 16p11.2 genes and n=37 for 22q11.2 genes) from 15633 genes and re-computed the PLSR 10000 times. The explained variance (R-squared) was used as test-statistics for the null distribution, and the p-value was calculated as the number of times the explained variance of the random gene-set exceeded the variance explained by 16p11.2 or 22q11.2 genes. A similar approach was performed for the individual gene correlations using median correlation as test-statistics.

### Data availability

Beta maps from all Connectome wide association studies (16p11.2 deletion and duplication, 22q11.2 deletion and duplication, ASD, SZ, and ADHD) performed in this study are available in the supplemental table 1 and on GitHub. Expression data used in this study are also available in supplemental table 4.

### Code availability

The processing scripts and custom analysis software used in this work are available in a publicly accessible GitHub repository with instructions on how to set up a similar computation environment and with examples of key visualizations in the paper: https://github.com/surchs/Neuropsychiatric_CNV_code_supplement

## Supporting information

Supplementary Material, Methods, and Results

## Supplementary Materials, Methods and Results

**Supplementary Material, Methods, and Results**
**Supplemental Material and Methods**
  Objectives and methods overview
  Samples
    16p11.2 cohort
    22q11.2 cohort
    Idiopathic ASD dataset
    Idiopathic schizophrenia
    Idiopathic ADHD
  Preprocessing
  Quality Control
  Aligning the gene expression maps from AHBA to the MIST64 functional parcellation
  Additional information on motion for each cohort after preprocessing
**Supplemental Results**
  Sensitivity analyses psychiatric diagnoses in 22q11.2 deletion
  Sensitivity analyses on age distribution in 16p11.2 deletion carriers
  Sensitivity analysis on the number of remaining frames in 16p11.2 deletion carriers
  Mirror effects of gene dosage in 16p11.2 CNV are present at the network level
  Effect of Schizophrenia on FC
  Effect of ASD on FC
  Effect of medication on FC alterations in autism
  Effect of ADHD on FC
  Effect of sex on FC alterations in SZ and ADHD
  Seed regions showing similarities between 16p11.2 deletion and Schizophrenia
  Seed regions showing similarities between 22q11.2 deletion and Schizophrenia
  Seed regions showing similarities between 16p11.2 deletion and Autism
  Seed regions showing significant similarity between 22q11.2 deletion and Autism
  Do the same seed regions contribute to the similarity between either 16p11.2 or 22q11.2 deletions and individuals with SZ and ASD?
  Regional similarities between the individual FC profiles of subjects with a psychiatric diagnosis and FC-signatures of 16p11.2 and 22q11.2 duplications
  Similarity between the individual FC profiles of subjects with a psychiatric diagnosis and the FC-signatures of the 16p11.2 and 22q11.2 deletions and duplications

## Supplemental figures and external tables

- Figure 1. Sensitivity analyses on psychiatric diagnoses in 22q11.2 deletion
- Figure 2. Sensitivity analyses on age distribution in 16p11.2 deletion carriers
- Figure 3. Mirror effects of gene dosage in 16p11.2 CNV are present at the network level
- Figure 4. Effects of ASD and SZ on FC before and after adjustment for global signal
- Figure 5. Seed regions showing similarities between 16p11.2 and 22q11.2 deletions, ASD and Schizophrenia
- Figure 6. Do the same seed regions contribute to the similarity between either 16p11.2 or 22q11.2 deletions and individuals with SZ and ASD?
- Figure 7. Regional similarities between the individual FC profiles of subjects with a psychiatric diagnosis and FC-signatures of 16p11.2 and 22q11.2 duplications
- Figure 8. Similarity between the individual FC profiles of subjects with a psychiatric diagnosis and the FC-signatures of the 16p11.2 and 22q11.2 deletions and duplications
- Table S1. CWAS beta estimates, ranking, region and networks labels.
- Table S2. Similarity of individuals with idiopathic psychiatric disorders with deletion FC-signatures
- Table S3. Association between similarity with deletion FC-signatures and symptom severity
- Table S4. Association between FC-signatures and spatial patterns of gene expression

## Author contributions

C.M., S.U., S.J., and P. B., designed the overall study and drafted the manuscript.

C.M. and S.U. processed the data and performed all imaging analyses.

P.O. preprocessed the SZ data and reviewed the manuscript.

C.S. performed the statistical analyses and drafted the manuscript.

K.K. and G.D. performed the gene expression analyses

A.L., E.D., PO, and G.H. contributed to interpretation of the data and reviewed the manuscript.

A.L. and L.K. provided the UCLA fMRI data.

A.E. contributed to interpretation of the data and reviewed the manuscript.

S.G., D.L., A.M., S.P., and E.S. provided the SZ data.

CE.B. provided the UCLA fMRI data, contributed to interpretation of the data and drafted the manuscript.

The Simons Variation in Individuals Project Consortium provided the 16p11.2 data.

All authors provided feedback on the manuscript.

