## Supplementary Material, Methods, and Results for "Neuropsychiatric mutations delineate functional brain connectivity dimensions contributing to autism and schizophrenia"

**Supplementary Material, Methods, and Results** 0

**Supplemental Material and Methods** 1

Objectives and methods overview 1

Samples 2

16p11.2 cohort 2

22q11.2 cohort 2

Idiopathic ASD dataset 3

Idiopathic schizophrenia 3

Idiopathic ADHD 4

Preprocessing 4

Quality Control 5

Aligning the gene expression maps from AHBA to the MIST64 functional parcellation 5

Additional information on motion for each cohort after preprocessing 6

**Supplemental Results** 6

Sensitivity analyses psychiatric diagnoses in 22q11.2 deletion 6

Sensitivity analyses on age distribution in 16p11.2 deletion carriers 7

Sensitivity analysis on the number of remaining frames in 16p11.2 deletion carriers 8

Mirror effects of gene dosage in 16p11.2 CNV are present at the network level 8

Effect of Schizophrenia on FC 10

Effect of ASD on FC 10

Effect of medication on FC alterations in autism 11

Effect of ADHD on FC 12

Effect of sex on FC alterations in SZ and ADHD 12

Seed regions showing similarities between 16p11.2 deletion and Schizophrenia 13

Seed regions showing similarities between 22q11.2 deletion and Schizophrenia 14

Seed regions showing similarities between 16p11.2 deletion and Autism 16

Seed regions showing significant similarity between 22q11.2 deletion and Autism 17

Do the same seed regions contribute to the similarity between either 16p11.2 or 22q11.2 deletions and individuals with SZ and ASD? 18

Regional similarities between the individual FC profiles of subjects with a psychiatric diagnosis and FC-signatures of 16p11.2 and 22q11.2 duplications 19

Similarity between the individual FC profiles of subjects with a psychiatric diagnosis and the FC-signatures of the 16p11.2 and 22q11.2 deletions and duplications 21

**References** 23

### Supplemental Material and Methods

#### Objectives and methods overview

| Aim | Objective | Method |
|---|---|---|
| AIM1.<br>Characterize the impact of gene dosage on connectivity for CNVs at the 16p11.2 and 22q11.2 genomic loci | 1.1. Describe the effect of gene dosage at the 16p11.2 and 22q11.2 genomic loci on global FC. | Connectome Wide Association Study (CWAS): Linear model contrasting CNV carriers with respective controls. |
| | 1.2 Test if deletions and duplications have mirror effects at the connection level. | |
| AIM2.<br>Test if FC-signatures of deletions represent dimensions observed in idiopathic ASD, SZ, or ADHD. | 2.1 Characterize the effect size of idiopathic Autism, Schizophrenia, and ADHD on FC. | Same as 1.1<br>CWAS: Linear model contrasting cases of each psychiatric groups with respective controls. |
| | 2.2 Test similarities between whole-brain connectomes of CNVs and idiopathic psychiatric conditions. | Pearson correlation between beta maps obtained from each CWAS (case-control contrast) and individual connectomes of either cases or controls of each group. |
| | 2.3 Investigate regions that contribute to similarities observed in 2.2 | The same method used in 2.2 applied to each of the 64 regional FC patterns. |
| | 2.4 Identify the relationship between regions identified in 2.3 and cognition | Pearson R computed in 2.3 was correlated to the cognitive measure of each individual |
| | 2.5 Associate the deletion FC-signatures with spatial patterns of genes expression within both loci | Partial Least Square Regression and correlation analyses between spatial patterns of gene expression and FC-signatures |

Table 1 Aims, objectives and methods

### Samples

#### 16p11.2 cohort

Imaging data of 16p11.2 CNV carriers and typically developing controls were acquired by the Simons variation in individuals project (VIP) consortium [1] across 2 sites. We excluded 44 individuals from the analysis due to insufficient quality of the imaging data (cf. Supplementary Methods, quality control). The final 16p11.2 sample includes 122 individuals. Over 90% of the deletion carriers and 69% of the duplication carriers met criteria for at least one clinical psychiatric diagnosis (Table 1**)**. Control subjects were recruited from the general population (extra-familial subjects), and had no major DSM-V diagnosis. The duplication group includes 2 individuals with a triplication.

#### 22q11.2 cohort

Imaging data of 22q11.2 CNV carriers and typically developing (TD) controls were acquired at the University of California, Los Angeles (UCLA). Patients were ascertained from the UCLA or Children's Hospital, Los Angeles Pediatric Genetics, Allergy/Immunology and/or Craniofacial Clinics. We excluded 16 individuals from the analysis due to insufficient quality of the imaging data (cf. Supplementary Methods, quality control). The final 22q11.2 sample includes 101 individuals. Demographically comparable TD comparison subjects were recruited from the same communities as patients via web-based advertisements and by posting flyers and brochures at local schools, pediatric clinics, and other community sites. Exclusion criteria for all study participants included significant neurological or medical conditions (unrelated to 22q11.2 mutation) that might affect brain structure, history of head injury with loss of consciousness, insufficient fluency in English, and/or substance or alcohol abuse or dependence within the past 6 months. The UCLA Institutional Review Board approved all study procedures and informed consent documents. Scanning was conducted on an identical 3 tesla Siemens Trio MRI scanner with a 12-channel head coil at the University of California at Los Angeles Brain Mapping Center or at the Center for Cognitive Neuroscience [2].

#### Idiopathic ASD dataset

The ABIDE dataset[3] is an aggregate sample of different studies including imaging and behavioural data for individuals with an ASD diagnosis and typically developing peers matched for age. Due to the small number of females in the ABIDE dataset, we excluded female individuals. To better account for biases in connectivity estimation due to differences in recording sites, subject age, and in scanner motion, we created age and motion-matched subsamples for each recording site in ABIDE of individuals that passed our quality control criteria. We then excluded recording sites with fewer than 20 individuals (10 ASD, 10 controls). Our final ABIDE sample thus includes 459 male individuals, 225 individuals with ASD and 234 healthy controls, from 10 recording sites.

#### Idiopathic schizophrenia

We used fMRI data retrospectively aggregated from 10 distinct sites and studies. Brain imaging multi-state data were obtained through either the SchizConnect and OpenfMRI data sharing platforms (http://schizconnect.org[4];https://openfmri.org[5]) or local scanning at the University of Montréal. All patients were diagnosed with SZ according to DSM-IV or DSM-V criteria, as a function of the time of study. Sites samples were obtained after subjects were selected in order to ensure even proportions of SZ patients and controls within each site (from N = 9 to N = 42 per group) and to reduce between-group differences with regards to gender ratio (74% vs. 75% males in patients and controls, respectively), age distribution (34 vs. 32 years old on average) and motion levels (averaged frame displacement: 0.16 vs. 0.14 mm). Such matching of SZ and controls subjects was achieved based on propensity scores. In total, we retained 242 SZ patients and 242 healthy controls in statistical analyses.

Depending on the study, positive and negative symptoms were assessed with either with the Positive and negative syndrome scale (PANSS,[6]) or the Scales for the assessment of positive/negative symptoms (SAPS/SANS,[7]). In order to allow for group analyses, SAPS/SANS scores were converted into PANSS scores using published regression-based equations[8].

#### Idiopathic ADHD

We used data provided by the ADHD-200 Consortium and The Neuro Bureau ADHD-200 Preprocessed repository (8 cohorts http://fcon_1000.projects.nitrc.org/indi/adhd200/ [9]). Data from seven sites were retained after exclusion of 184 individuals. We included in our study a total of 763 subjects, 289 patients diagnosed with ADHD and 474 healthy controls.

This database provided child and adolescent data. Scores related to ADHD symptoms were measured using Conner's Parent Rating Scale-Revised, Long Version (CPRS-LV [10]).

### Preprocessing

All datasets were preprocessed using the same parameters with the same Neuroimaging Analysis Kit (NIAK) version 0.12.4, an Octave-based open-source processing and analysis pipeline [11]. The first four volumes of each rs-fMRI time series were discarded to allow for magnetization to reach a steady state. Each data set was corrected for differences in slice acquisition time. Head motion parameters were estimated by spatially re-aligning individual timepoints with the median volume in the time series. This reference median volume was then aligned with the individual anatomical T1 image, which in turn was co-registered onto the MNI152 template space using an initial affine transformation, followed by a nonlinear transformation. Finally, each individual timepoint was mapped to the MNI space [12] using the combined spatial transformations. Slow frequency drifts were modelled on the entire time series as discrete cosine basis functions with a 0.01 Hz high-pass cut-off. Timepoints with excessive in-scanner motion (greater than 0.5 mm framewise displacement) were then censored from the time series by removing the affected timepoint as well as the preceding and following two timepoints [13]. Nuisance covariates were regressed from the remaining time series: the previously estimated slow time drifts, the average signals in conservative masks of the white matter and lateral ventricles, and the first principal components (95% energy) of the estimated six rigid-body motion parameters and their squares. Data were then spatially smoothed with a 3D Gaussian kernel (FWHM = 6mm).

#### Quality Control

Preprocessed data were visually controlled for quality of the co-registration, head motion, and related artefacts by one rater. Not all six datasets were examined by the same raters, yet all raters followed the same standardized quality-control procedure [14]. If there was co-registration failure of either the functional image to individual T1 or individual T1 to MNI template registration, we attempted a manual fix by changing the parameters of the preprocessing pipeline. Individuals were excluded from the analysis if co-registration errors could not be fixed. Individuals were also excluded from the analysis if the average framewise displacement after motion censoring exceeded 0.5 mm or if fewer than 40 time frames remained (Supplementary Materials and Methods).

#### Aligning the gene expression maps from AHBA to the MIST64 functional parcellation

To investigate the transcriptomic relationship of altered FC in each deletion, we aligned publicly-available atlas of gene expression in the adult human cortex from the Allen Human Brain Atlas (AHBA) dataset [15] to the MIST64 brain parcellation following previously published guidelines for probe-to-gene mappings and intensity-based filtering [16] and adapting the abagen toolbox [17]. We normalized expression values within each brain sample across genes for each of the 6 donors and then for each gene across samples for each donor using a scaled robust sigmoid normalization [16]. We computed the mean of the normalized values of all samples encompassed within each functional region of the MIST64. This was performed for each donor and then averaged across donors. A leave-one-donor out sensitivity analysis generated 6 expression maps. The principal components of these 6 expression maps were highly correlated (average Pearson correlation of 0.993). The same high correlation was observed for the differential Stability score (average Pearson correlation of 0.987).

Normalized gene expression value was available for each of the 15663 genes and for each of the 64 functional brain regions.

##### Additional information on motion for each cohort after preprocessing

| | ADHD | CON | SZ | CON | ASD | CON | 16PDEL | 16PDUP | 16PCON | 22QDEL | 22QDUP | 22QCON |
|---|---|---|---|---|---|---|---|---|---|---|---|---|
| Mean (frames) | 160 | 159 | 152 | 166 | 139 | 153 | 83 | 83 | 85 | 110 | 121 | 126 |
| Sd (frames) | 53 | 59 | 50 | 49 | 52 | 49 | 18 | 23 | 20 | 42 | 34 | 30 |
| Mean (% scrubbed) | 0.13 | 0.25 | * | * | 0.24 | 0.16 | 0.26 | 0.25 | 0.23 | 0.25 | 0.18 | 0.14 |
| Sd (% scrubbed) | 0.15 | 0.26 | * | * | 0.23 | 0.18 | 0.15 | 0.2 | 0.16 | 0.28 | 0.22 | 0.2 |

Table 2: Mean (frames): mean number of frames remaining after scrubbing per individual; Mean (% scrubbed): Mean percentage of frames excluded per individual; SD: Standard Deviation; 16PDEL: 16p11.2 deletion; 16PCON: 16p11.2 control; 16PDUP: 16p11.2 duplication; 22QDEL: 22QCON: 22q11.2 control; 22q11.2 deletion; 22QDUP: 22q11.2 duplication. Sensitivity analyses on the number of remaining frames in 16p11.2 deletion carriers are available below.

#### Supplemental Results

##### Sensitivity analyses on psychiatric diagnoses in 22q11.2 deletion

We compared the FC signatures of 22q11.2 deletion carriers with and without ASD. Both FC signatures showed the same global underconnectivity and were strongly correlated with the one presented in the manuscript despite the smaller sample size (r=0.83 to 0.90). The same sensitivity analysis also showed that ADHD diagnosis had no detectable influence on the 22q11.2 deletion FC signature. We performed the similarity analyses between 22q11.2 deletion profiles and idiopathic conditions after exclusion of 22q11.2 subjects with either ASD or ADHD. The same regions were driving similarities with 22q11.2 FC profiles irrespective of diagnoses.

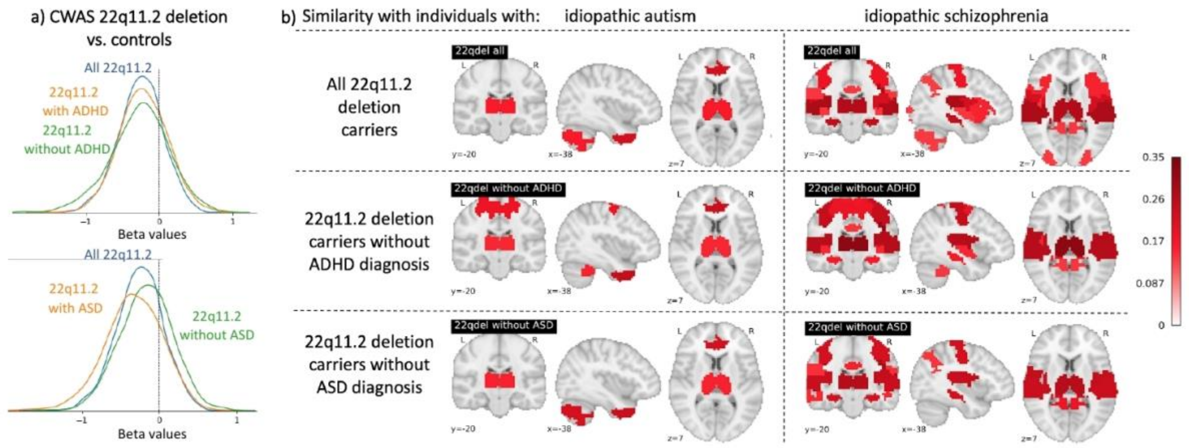


Figure 1: Legend: a) Density plots show the distribution of beta coefficients of CWAS contrasting controls and subgroups of 22q11.2 deletion (with and without diagnoses, in orange and green respectively). The full 22q11.2 sample is in blue. Under-connectivity is present in all 22q11.2 subgroups irrespective of diagnosis. b) Similarities between 22q11.2 regional FC-signature and individuals with ASD and SZ. The same regions showed similarities in all 22q11.2 subgroups irrespective of psychiatric diagnosis, despite the 2-fold decrease in sample size. Maps are thresholded maps (FDR). Color scale represents the similarity effect size (Mann Whitney, rank biserial correlation).

#### Sensitivity analyses on age distribution in 16p11.2 deletion carriers

We performed a sensitivity analysis after excluding older controls to obtain identical age distributions (mean 12.7 and 13.0 years respectively) in the 16p11.2 deletions and control groups. The CWAS performed before and after excluding adults provides the same results albeit with decreased power: The 2 beta maps are highly correlated (r=0.967) and their distribution (b) overlaps perfectly.

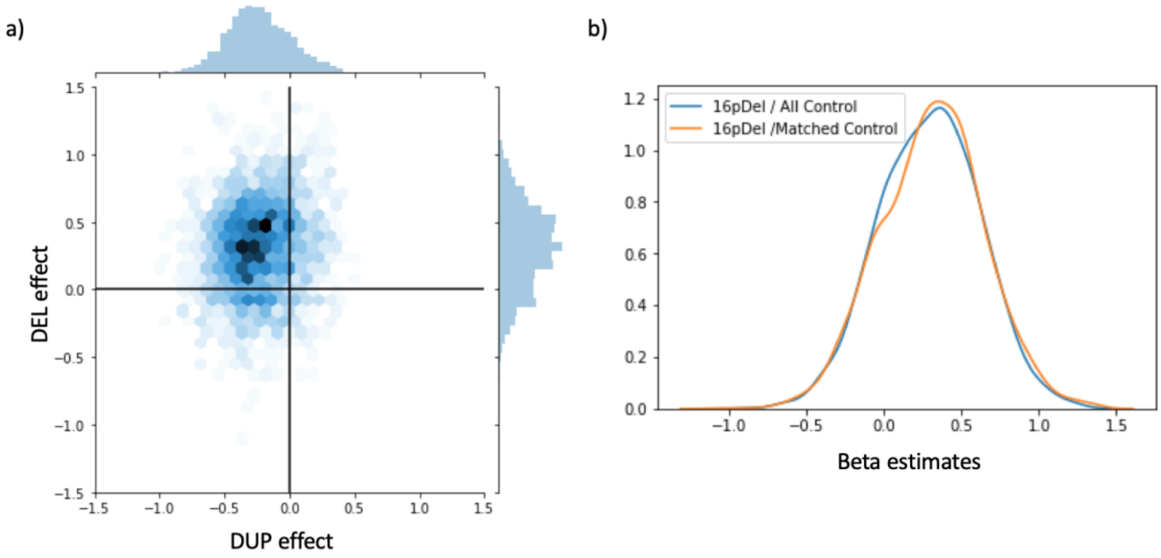


Figure 2: Legend: (a): Scatterplot (*hexagonal plot*), showing estimates (beta values) from connectome-wide association studies (CWAS) performed between 16p11.2 CNVs and their respective controls excluding adults to obtain a matched age distribution. The scatterplot is identical to figure 1.a demonstrating that age distribution does not affect our results.

#### Sensitivity analysis on the number of remaining frames in 16p11.2 deletion carriers

We performed a sensitivity analysis by excluding all individuals with less than 60 frames (n=4 deletion carriers, 4 duplication carriers, 10 controls). The CWAS results were very close to those obtained with all participants. FC-signatures before and after exclusion showed an r=0.94 and r=0.89 for 16p11.2 deletions and duplications respectively.

#### Mirror effects of gene dosage in 16p11.2 CNV are present at the network level

We investigated how mirror effects of gene dosage at the individual network level follow the organization of the brain into canonical resting-state networks.

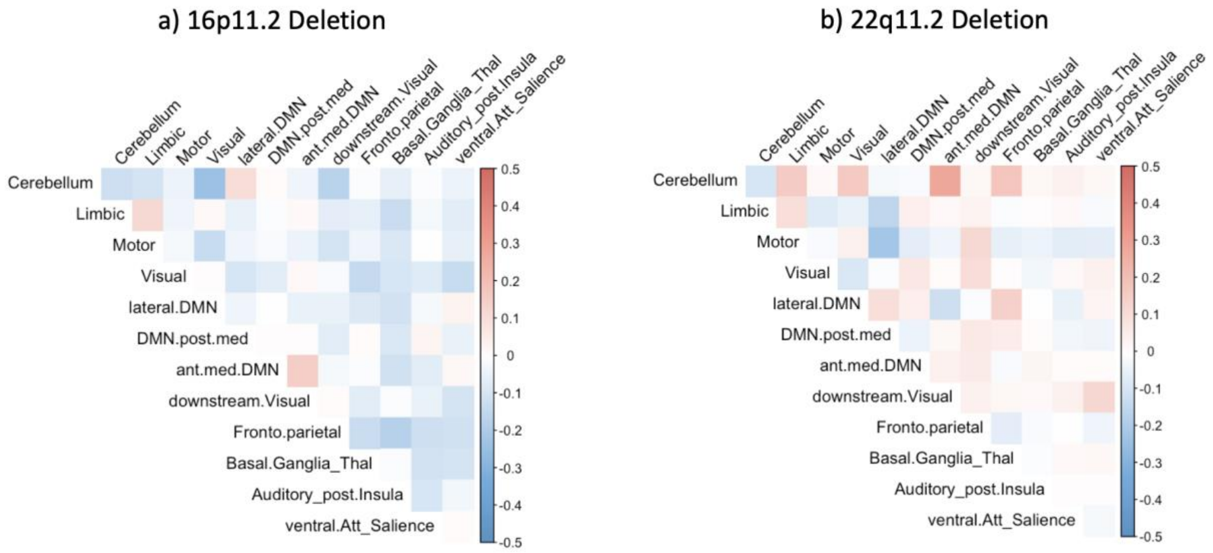


Figure 3: Legend: Gene dosage effects at the connection level are shown broken down into the 12 corresponding functional brain networks (Supplementary Table S1.10) for the 16p11.2 (a) and 22q11.2 (b) genomic loci. Red colors reflect effects in the same direction for deletions and duplications. Blue colors reflect mirror effects for deletions and duplications (i.e. an increase in FC is associated with one, but a decrease is associated with the other). For the 16p11.2 CNVs, networks exhibited more mirror connectivity alterations than would be expected by chance (p = 0.006, two-sided). By contrast, 22q11.2 CNVs showed an equal number of connections with mirror and uni-directional alterations across networks.

#### Effect of Schizophrenia on FC

Idiopathic SZ showed overall underconnectivity affecting 835 connections, with the strongest alterations affecting the frontoparietal, auditory, ventral attention and limbic networks (dorsal anterior cingulate cortex, dorsal anterior and posterior insula, ventral posterior insula, and inferior marginal sulcus) (Figures 3a-b, Supplementary Tables S1.7 and S1.9, Supplementary Figure 4). Over-connectivity was restricted to 24 connections (cerebellar-motor and thalamic-auditory, Figure 3c-d), in line with previous reports [18–20] (859 connections survived FDR correction in total, $q < 0.05$, p ranging from 0.02 to 1e-12).

#### Effect of ASD on FC

Underconnectivity in ASD was driven by seed regions in the limbic, lateral DMN, and cerebellar networks and over-connectivity was restricted to the thalamus connectivity profile, with the medial somatomotor network (MOTnet_m), in accordance with previous reports [3,21,22] (temporal pole, anterior middle temporal gyri, lateral fusiform gyrus, anterior middle frontal gyrus, and cerebellum VIII-ab) (Figures 3, Supplementary Tables S1.5 and S1.9, Supplementary Figure 4). Seventy-five connections (73 under and 2 overconnected) survived FDR correction ($q < 0.05$), with p ranging from 0.01 to 1e-5.

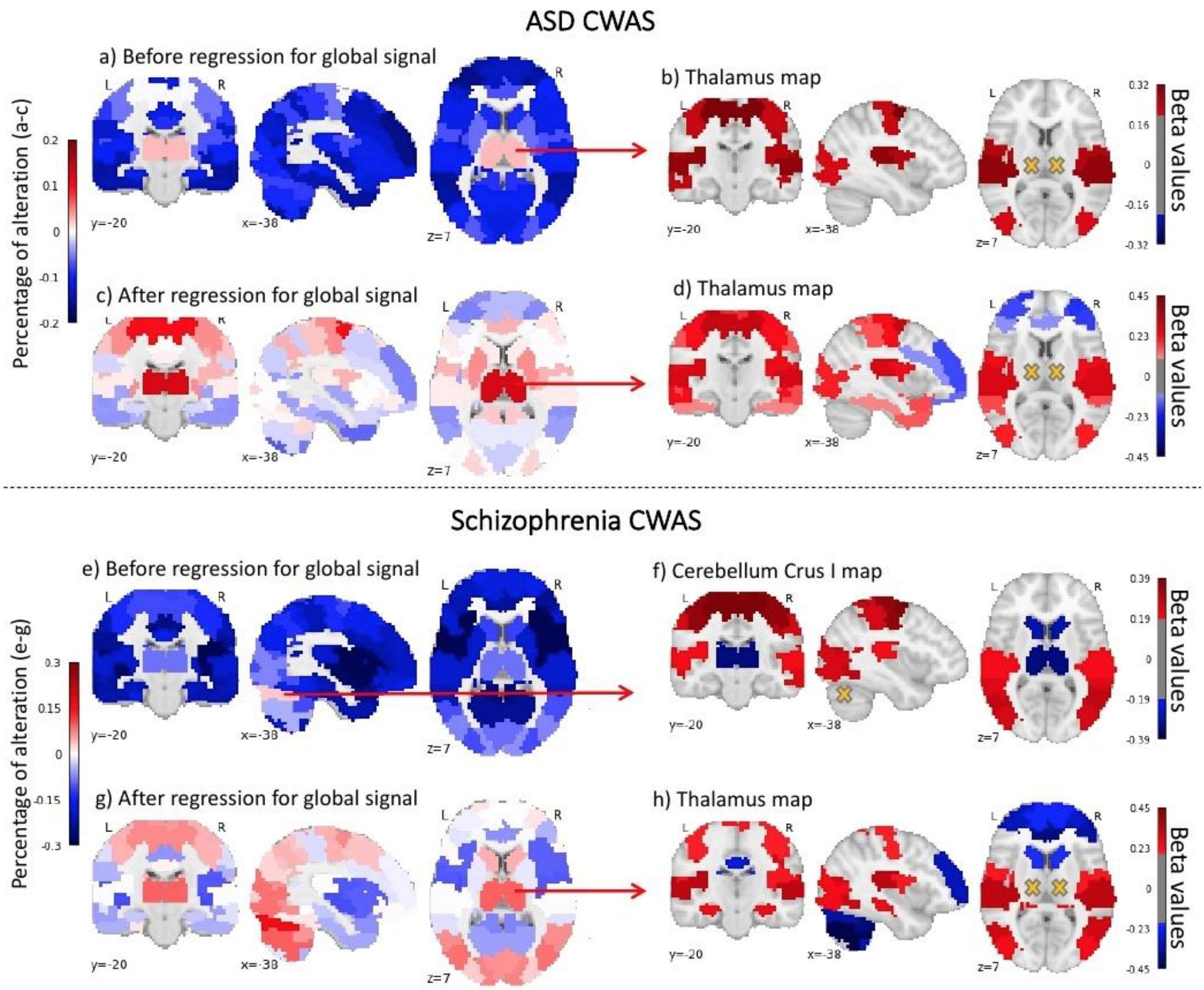


Figure 4: Effects of ASD and SZ on FC before and after adjustment for the global signal. The adjustment removes the mean shift (global underconnectivity observed in ASD and SZ). The alterations of each region and network relative to one another remain similar before and after adjustment and both FC-signatures (beta maps) are highly correlated (r=0.98). This also demonstrates that ASD and SZ are associated with a mean shift in connectivity as well as a reorganisation of networks relative to one another.

#### Effect of medication on FC alterations in autism

A sensitivity analysis testing the effect of medication was performed with excluding 55 individuals with autism and one control from the analysis. Connectome-wide analysis on the non-medicated subgroup showed overall underconnectivity. Underconnectivity was significant (FDR) in 28 connections. The beta map of this analysis was highly correlated (r= 0.96) with the initial CWAS performed on the full sample including ASD subjects with medication.

#### Effect of ADHD on FC

For ADHD, none of the individual connections survived FDR correction (nominally significant alterations were observed in the posterior middle temporal and lateral occipitotemporal gyri, posterior medial visual network, the cerebellum-VI and the left intraparietal sulcus) (Supplementary Tables S1.8 and S1.9).

#### Effect of sex on FC alterations in SZ and ADHD

A sensitivity analysis was performed on 179 male participants with schizophrenia and 189 male controls. Males with idiopathic SZ also show overall underconnectivity with 472 connections surviving FDR. The beta map of this analysis excluding females was highly correlated (r= 0.95) with the initial CWAS performed on the full sample.

The same analysis was performed in the ADHD sample. The beta map of the analysis excluding females was highly correlated (r= 0.93) with the initial CWAS performed on the full ADHD sample.

Seed regions showing similarities between 16p11.2 deletion and Schizophrenia

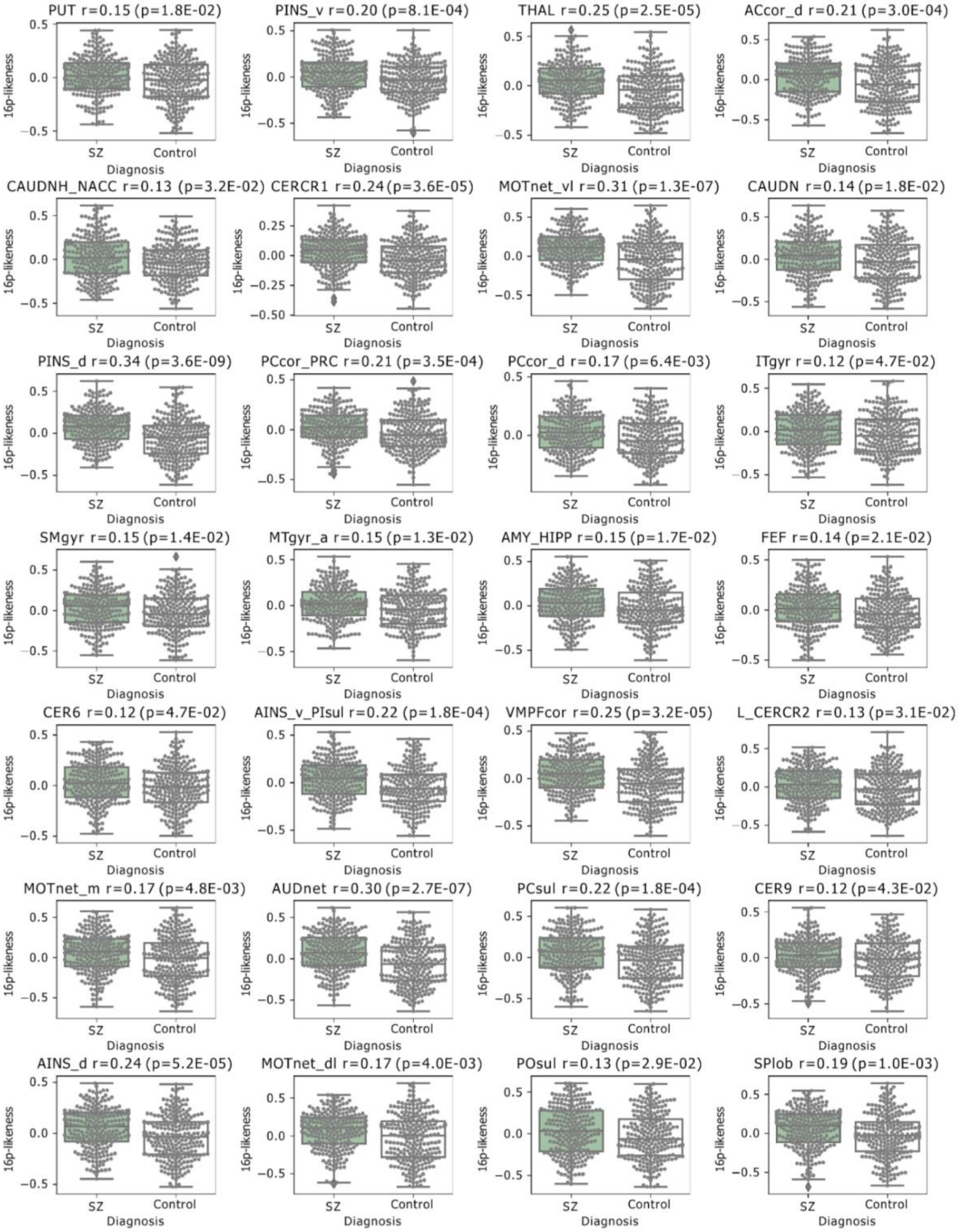


Figure 5a: Legend: Similarities between regional FC of 16p11.2 deletions and individuals with idiopathic SZ compared to their respective controls. Grey dots correspond to individuals in either the SZ or control group. Boxplots for individuals with SZ (green) and their respective controls (white)

illustrate the observed group differences in similarity values. Similarity values (Pearson's R, Grey dots) were derived by computing Pearson's correlations between the regional FC-signatures. Differences between SZ or controls are tested using a Mann-Whitney U test. We reported significant group differences after FDR correction accounting for the 64 regions ($q < 0.05$).

#### Seed regions showing similarities between 22q11.2 deletion and Schizophrenia

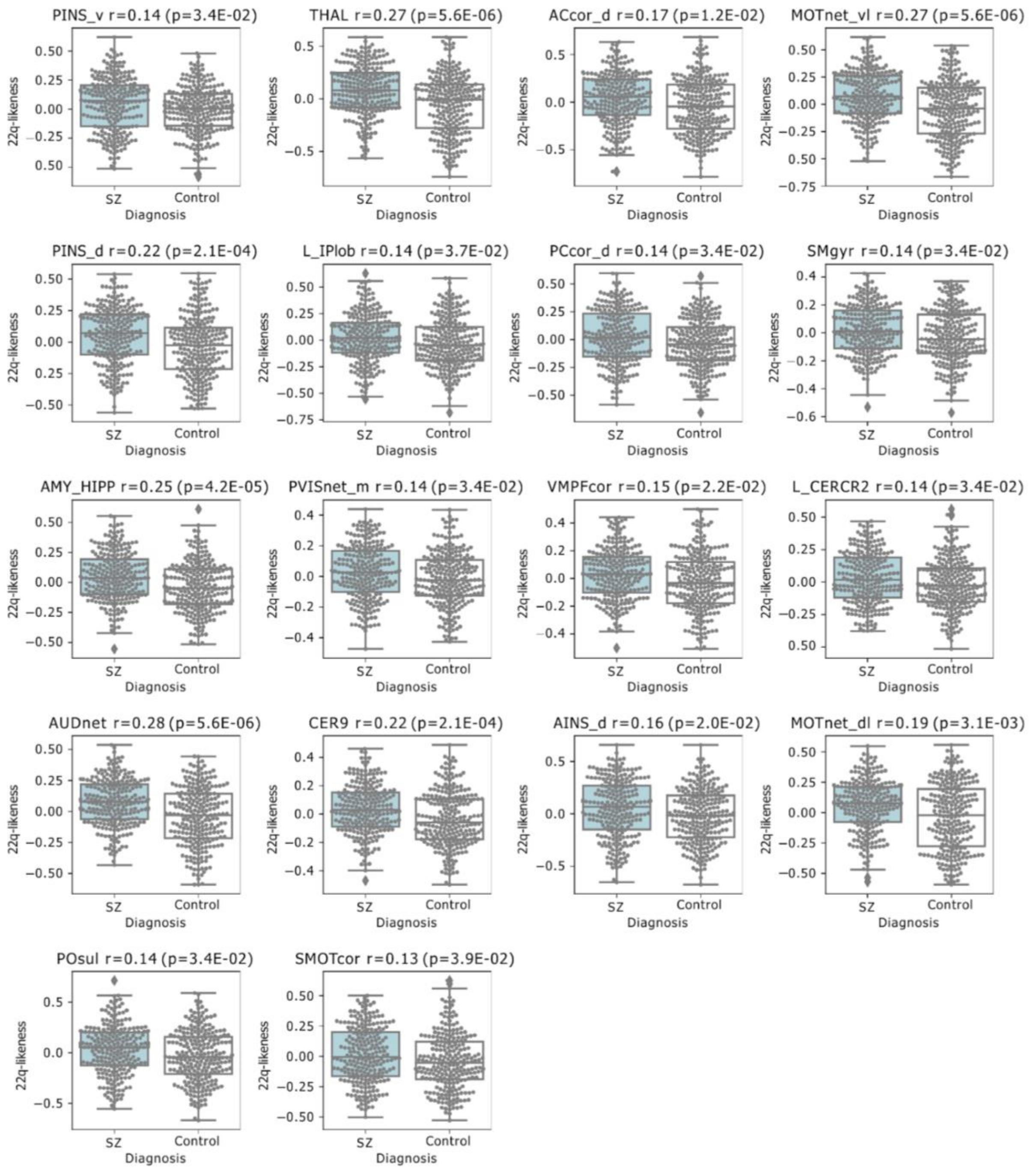

Figure 5b: Legend: Similarities between regional FC of 22q11.2 deletions and individuals with SZ compared to controls. Grey dots correspond to individuals in either the SZ or control group. Box plots for individuals with SZ (blue) and their respective controls (white) illustrate the observed group differences in similarity values.

Similarity values (Pearson's R, Grey dots) were derived by computing Pearson's correlations between the regional FC-signatures. Differences between SZ or controls are tested using a Mann-Whitney U test. We reported significant group differences after FDR correction accounting for the 64 regions ($q < 0.05$).

#### Seed regions showing similarities between 16p11.2 deletion and Autism

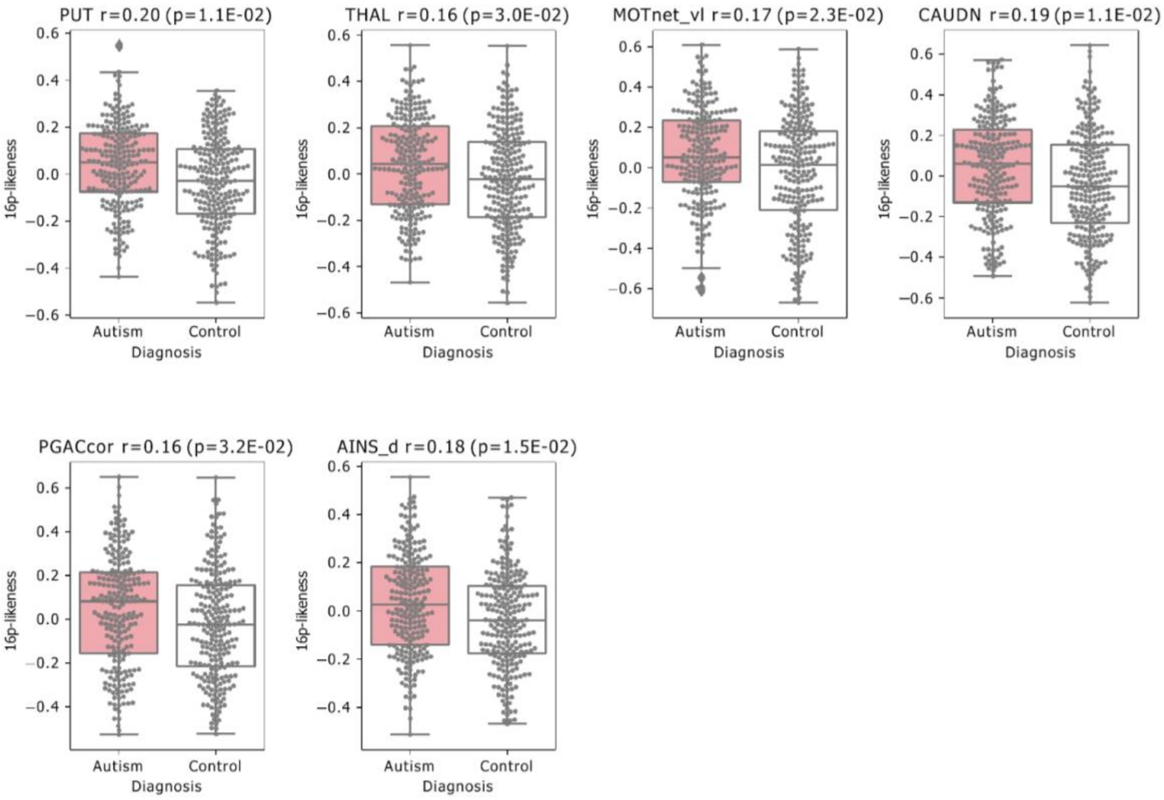


Figure 5.c Legend: Similarities between regional FC of 16p11.2 deletions and individuals with ASD compared to controls are shown. Grey dots correspond to individuals in either the ASD or control group. Box plots for individuals with ASD (red) and their respective controls (white) illustrate the observed group differences in similarity values. Similarity values (Pearson’s R, Grey dots) were derived by computing Pearson’s correlations between the regional FC-signatures. Differences between SZ or controls are tested using a Mann-Whitney U test. We reported significant group differences after FDR correction accounting for the 64 regions ($q < 0.05$).

#### Seed regions showing significant similarity between 22q11.2 deletion and Autism

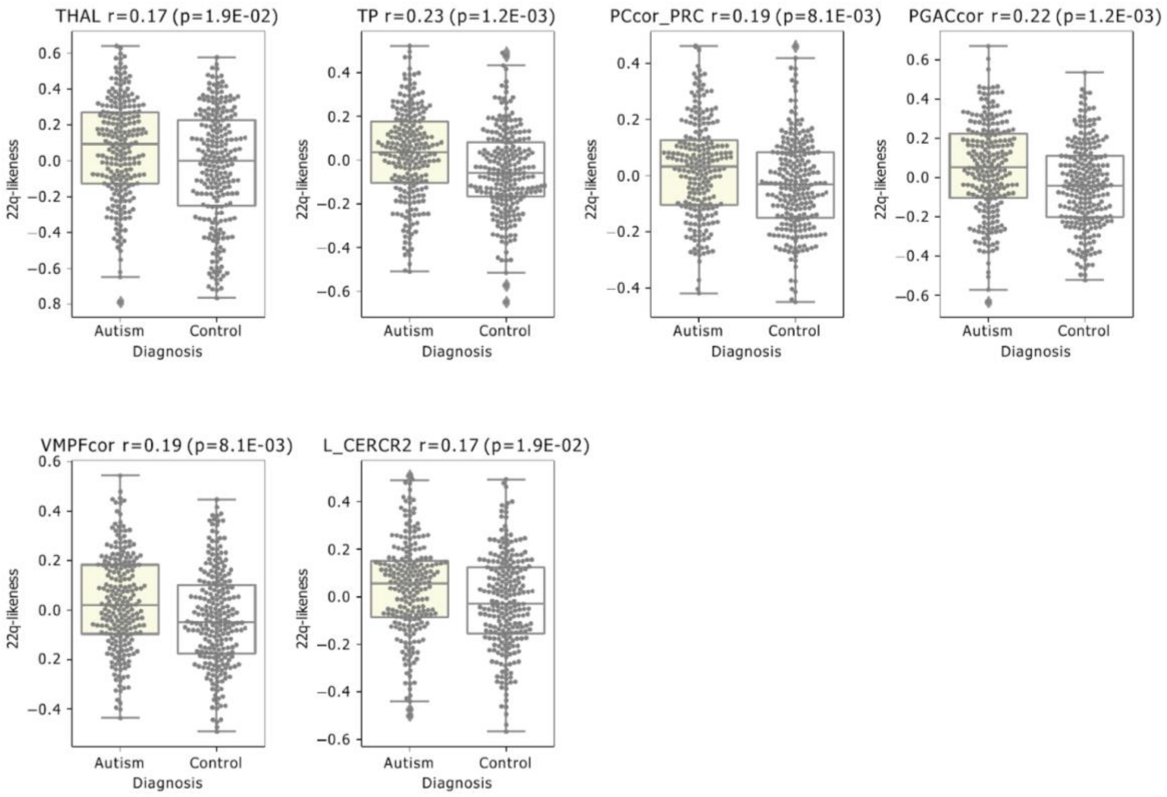


Figure 5.d Legend: Similarities between regional FC of 22q11.2 deletions and individuals with ASD compared to controls are shown. Grey dots correspond to individuals in either the ASD or control group. Box plots for individuals with ASD (yellow) and their respective controls (white) illustrate the observed group differences in similarity values. Similarity values (Pearson's R, Grey dots) were derived by computing Pearson's correlations between the regional FC-signatures. Differences between SZ or controls are tested using a Mann-Whitney U test. We reported significant group differences after FDR correction accounting for the 64 regions ($q < 0.05$).

#### Do the same seed regions contribute to the similarity between either 16p11.2 or 22q11.2 deletions and individuals with SZ and ASD?

Many of the same seed regions contribute to the similarity between either 16p11.2 or 22q11.2 deletions and individuals with SZ (a). This is less the case for autism.

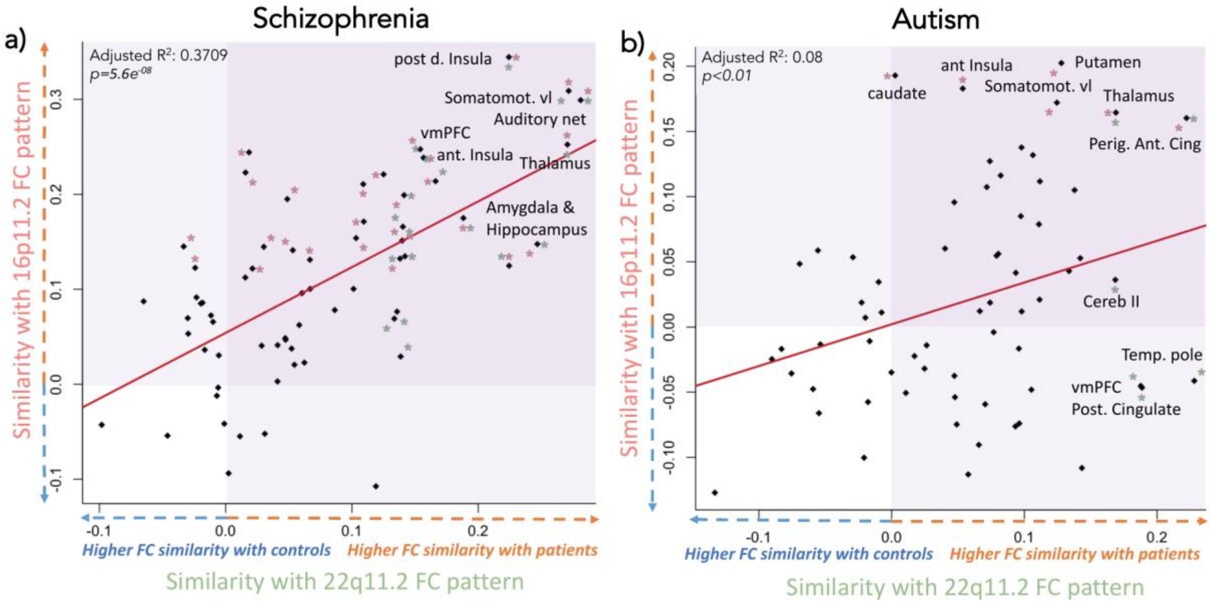


Figure 6. Legend: The horizontal axes show the Mann Whitney effect size (rank biserial correlation) representing the connectivity similarity between the 22q11.2 deletion FC-signature and individuals with idiopathic SZ (a) or ASD (b) compared to controls. The vertical axes show values (biserial correlation) representing the connectivity similarity with the 16p11.2 deletion FC-signature and individuals with idiopathic ASD (a) or SZ (b) compared to controls. Positive values (orange arrows) on the horizontal and vertical axes represent higher connectivity similarity between the deletion FC-signatures and individuals with SZ (a) or ASD (b) compared to their respective controls. Negative values (blue arrows) represent a higher similarity with control individuals. Seed-based connectivity signatures that show significant (FDR corrected) connectivity similarity with deletions are coloured (pink stars for 16p11.2 and green stars for 22q11.2).

#### Regional similarities between the individual FC profiles of subjects with a psychiatric diagnosis and FC-signatures of 16p11.2 and 22q11.2 duplications

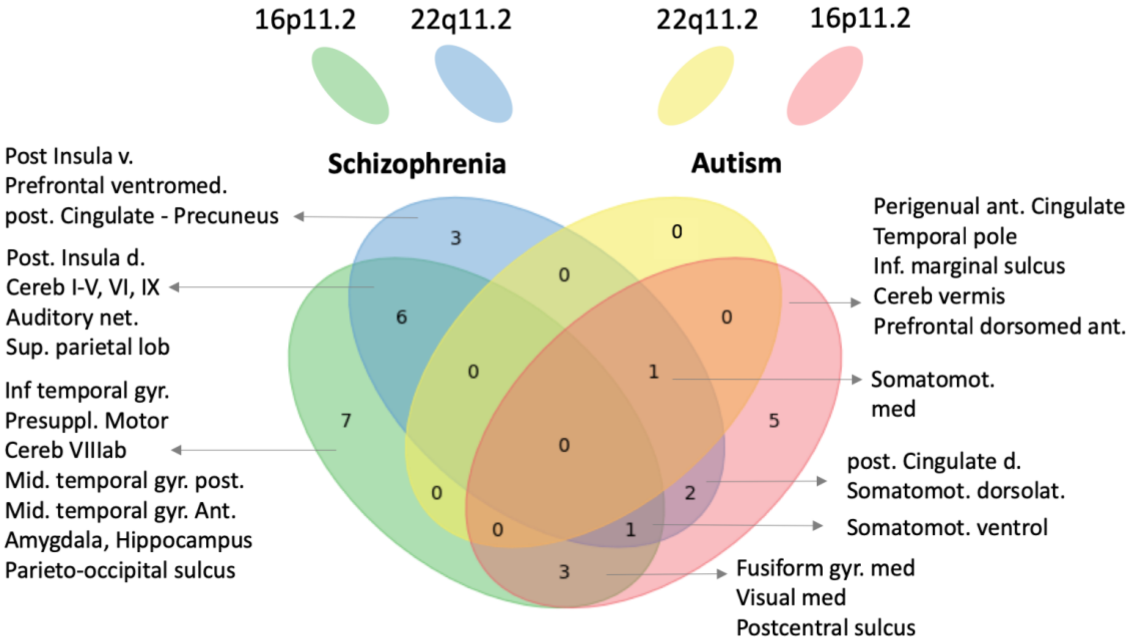


Figure 7. Legend: The FC-signatures of both duplications (group level) were decomposed into 64 seed-regions. Deletion FC-signatures are correlated to the individual connectomes of subjects with either ASD, SZ or control subjects. Of note, the correlation results in the mean centring of all region-based FC-signatures. Significantly increased similarities with either ASD and SZ are presented on the right and the left side of the diagram, respectively.

Some of these regions overlap with those identified in the same analysis using FC-signatures of both deletions: SZ individuals show similarities with both 16p11.2 deletion and duplication for 2 regions (Inferior and middle temporal gyrus)

SZ individuals show similarities with both 22q11.2 duplication and deletion for 6 regions (ventral/dorsal posterior insula, ventromedial prefrontal cortex, dorsolateral somatomotor cortex, cerebellum IX, auditory network) and with 16p11.2 duplication and deletions for 9 regions (parietal occipital sulcus, amygdala/ hippocampus, inf & middle gyrus anterior and posterior temporal gyrus, superior parietal lobule, cerebellum VI, posterior insula, dorsal prefrontal ventromedial cortex). ASD individuals show similarities with both 16p11.2 deletion and duplication for 2 regions perigenual anterior cingulate cortex, ventrolateral somatomotor cortex.

For 11 regions, controls rather than individuals with ASD or SZ showed increased similarities with duplications. This increased similarity with controls was never observed for deletion FC-signatures.

Similarity between the individual FC profiles of subjects with a psychiatric diagnosis and the FC-signatures of the 16p11.2 and 22q11.2 deletions and duplications

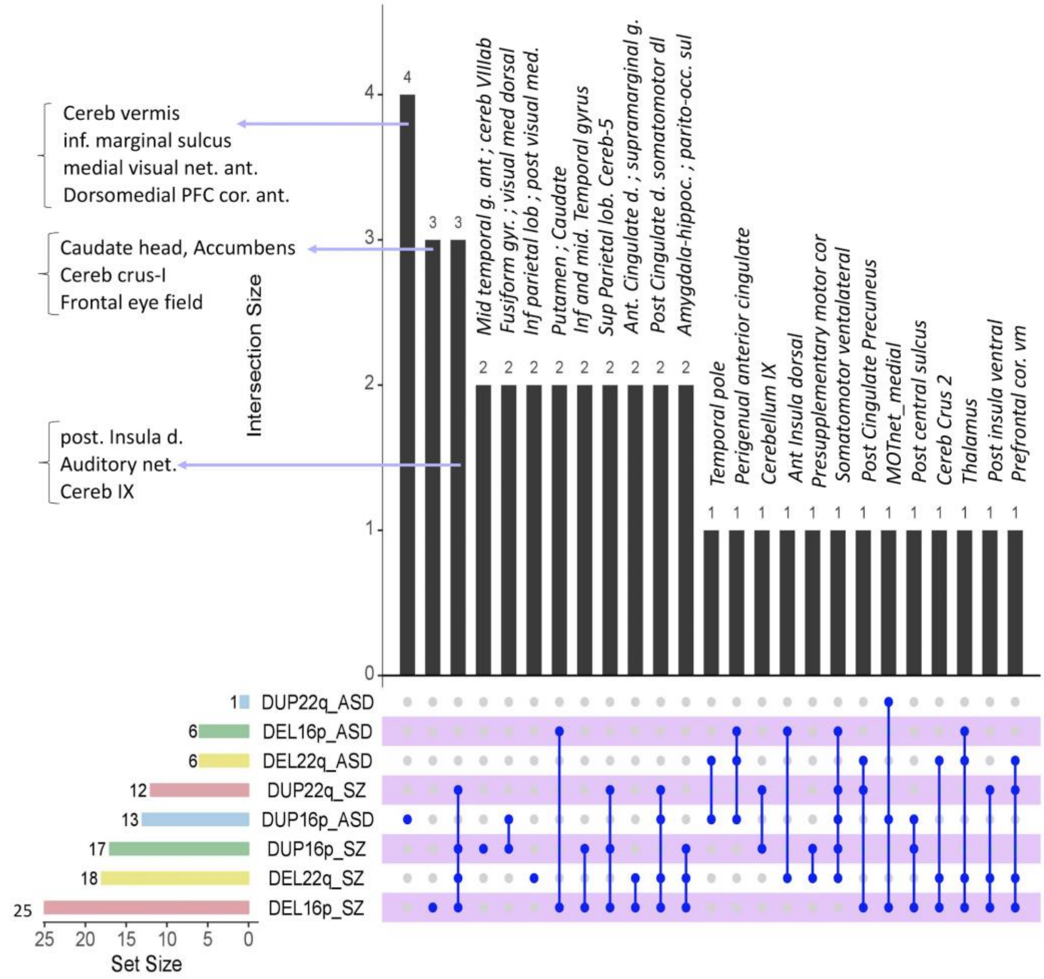


Figure 8. Legend: Because a Venn diagram showing overlapping brain regions across 8 comparisons (conditions, dels and dups) is impractical, we used the UpSet R package 23 to combine results from both Venn-Diagram (presented in Figure 4 (deletion) and in Supplementary Figure 6 (duplication)).

The dark blue lines and points show regions contributing to several similarities.

The 8 rows represent the 8 comparisons between the 4 CNVs and idiopathic ASD or SZ.

For example, the 23rd vertical black bar represents the thalamus which contributes to similarities across 4 comparisons (the 4 connected dark blue dots, representing the 16p11.2 and 22q11.2 deletion-FC-similarity with either ASD or SZ).

Horizontal bars on the bottom left represent the number of seed regions showing significant similarities for each comparison (eg. 6 regions showed similarity between 16p11.2 deletion-FC-signatures and ASD-FC-profiles). Y-axis: number of regions showing similarities between CNV carriers and idiopathic SZ or ASD.
